# Structural and Functional Insights into the Action Mode of A Mitochondrial AAA+ Disaggregase CLPB

**DOI:** 10.1101/2022.03.10.483744

**Authors:** Damu Wu, Yan Liu, Yuhao Dai, Guopeng Wang, Guoliang Lu, Yan Chen, Ningning Li, Jinzhong Lin, Ning Gao

**Affiliations:** State Key Laboratory of Membrane Biology, Peking-Tsinghua Joint Center for Life Sciences, School of Life Sciences, Peking University, Beijing 100871, China.; State Key Laboratory of Genetic Engineering, School of Life Sciences, Zhongshan Hospital, Fudan University, Shanghai 200438, China; Academy of Advanced Interdisciplinary Studies, Peking University, Beijing 100871, China.

## Abstract

The human AAA+ ATPase CLPB (SKD3) is a protein disaggregase in the mitochondrial intermembrane space and functions to promote the solubilization of various mitochondrial proteins. CLPB deficiency by mutations is associated with a few human diseases with neutropenia and neurological disorders. Unlike canonical AAA+ proteins, CLPB contains a unique ankyrin repeat domain (ANK) at its N-terminus. The mechanism of CLPB functions as a disaggregase and the role of its ANK domain are currently unclear. Herein, we report a comprehensive structural characterization of human CLPB in both the apo- and substrate-bound states. CLPB assembles into homo- tetradecamers in apo-state and is remodeled into homo-dodecamers upon binding to substrates. Conserved pore- loops on the ATPase domains form a spiral staircase to grip and translocate the substrate in a step-size of two amino acid residues. The ANK domain is not only responsible for maintaining the higher-order assembly but also essential for the disaggregase activity. Interactome analysis suggests that the ANK domain may directly interact with a variety of mitochondrial substrates. These results reveal unique properties of CLPB as a general disaggregase in mitochondria and highlight its potential as a target for the treatment of various mitochondria-related diseases.

## Introduction

Protein misfolding and aberrant aggregation are devastating to many fundamental functions of the cell and failures to remediate them are closed related to many human diseases (Eisele et al., 2015; Soto and Pritzkow, 2018). To maintain a healthy proteome, cells have evolved multiple dedicated systems, one of which is the HSP100 chaperone family (Erzberger and Berger, 2006; Kirstein et al., 2009; Puchades et al., 2020). As a subfamily of the AAA+ ATPase, HSP100 proteins generally contain an N-terminal domain (NTD), one or two ATPase domains (or nucleotide-binding domain, NBD), and usually function in hexameric forms. Taking yeast Hsp104 as an example, the NTD is involved in substrate binding, while the NBD1 and NBD2 bind and hydrolyze ATP to facilitate power substrate unfolding and translocation (Grimminger-Marquardt and Lashuel, 2010). Similar to many other AAA+ ATPases, Hsp104 unfolds and transports substrates through its central pore by a ratchet-like motion of the highly conserved pore-loops (PL) on the ATPase domains (Gates et al., 2017). The ATP-hydrolysis cycle dependent conformational change of each subunit results in both inter- and intra-subunit structural remodeling, which collectively lead to the threading and unidirectional movement of the peptide within the central channel (Gates et al., 2017).

Recently, a new type of HSP100 family proteins, CLPB (also known as SKD3) was reported to act as a protein disaggregase in the intermembrane space (IMS) of mitochondria (Cupo and Shorter, 2020; Mroz et al., 2020; Thevarajan et al., 2020; Wortmann et al., 2015). The N-terminus of CLPB has a mitochondrial targeting signal (MTS), followed by an ankyrin repeat (ANK) domain, and ends with a C-terminal NBD (Fig. 1A). There is a short hydrophobic stretch between the MTS and the first ankyrin motif, as well as a long linker helix (LH) between ANK domain and NBD (Fig. 1A). The MTS is cleaved by mitochondrial processing peptidase (MPP), followed by a second cleavage by presenilin-associated rhomboid-like (PARL) protease to remove additional hydrophobic residues from the N-terminus (Saita et al., 2017; Thevarajan et al., 2020). The ANK domain is a unique feature of CLPB compared to other AAA+ ATPases, and the removal of ANK domain disrupts the disaggregase activity of CLPB (Cupo and Shorter, 2020).

**Figure 1.**
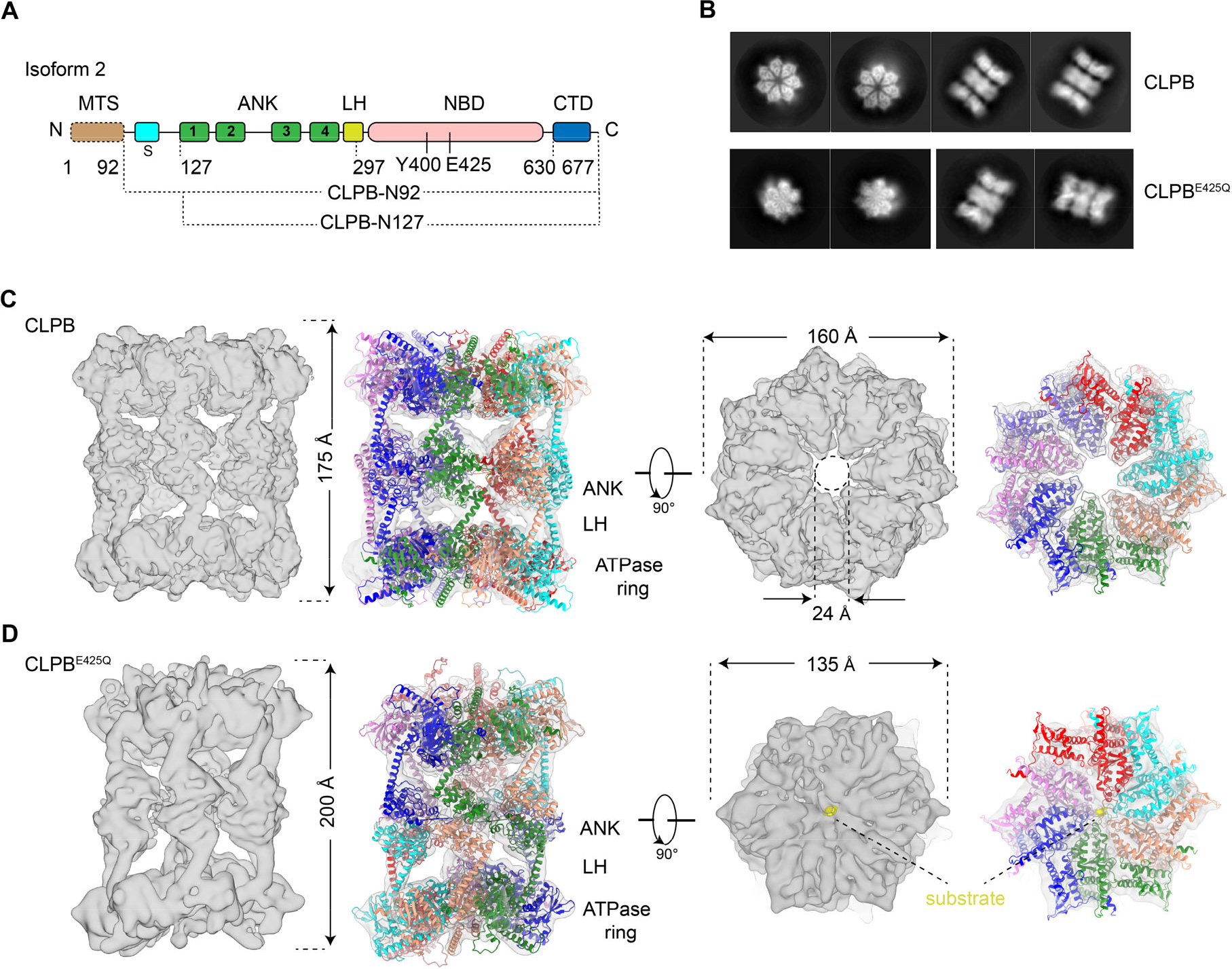
Cryo-EM structures of CLPB double-heptamers in the apo-state and CLPB^E425Q^ double-hexamers in the substrate-bound state. **A** Domain organization of *H. sapiens* CLPB. CLPB is composed of an MTS, a short hydrophobic stretch (S), a linker helix (LH), four ankyrin-repeat (ANK) motifs, an NBD, and a CTD. **B** Representative 2D classification averages of CLPB and CLPB^E425Q^ datasets. **C**, **D** The density maps of the double-heptamer (C) and double-hexamer (D), superimposed with the models of CLPB. The density maps are shown in the side and top (ATPase ring) views. The higher-order oligomer is mediated by ANK domains. LH, linker helix. The substrate was labelled as yellow.

Dysfunction of CLPB by mutations are associated with several human diseases, such as the 3-methylglutaconic aciduria (3-MGA) (Capo-Chichi et al., 2015; Kanabus et al., 2015; Kiykim et al., 2016; Saunders et al., 2015; Wortmann et al., 2015). A common disorder of the 3-MGA patients is the increased urinary 3-methylglutaric acid excretion, often with varying degree of microcephaly, small birth weight, neutropenia, severe encephalopathy, intellectual disability, movement disorder and cataracts (Capo-Chichi et al., 2015; Kanabus et al., 2015; Kiykim et al., 2016; Saunders et al., 2015; Wortmann et al., 2015). Moreover, heterozygous missense variants of CLPB were also identified in patients with severe congenital neutropenia (SCN), and these variants were found to disrupt granulocyte differentiation of human hematopoietic progenitors (Warren et al., 2022). Many of these disease-related mutations have been shown to impair the disaggregase activity of CLPB, such as T268M, R408G, R475Q, A591V, R650P in 3-MGA and N499K, E557K, R561G, R620C in SCN (Cupo and Shorter, 2020; Warren et al., 2022). At the cellular level, CLPB is important in maintaining normal cristae structure of mitochondria (Chen et al., 2019b). CLPB interacts with HAX1, an anti-apoptotic factor of BCL-2 family, to promote cell survival (Chen et al., 2019b; Wortmann et al., 2015). Upregulated cellular level of CLPB in acute myeloid leukemia (AML) cells was found to mediate the resistance to BCL-2 inhibitor venetoclax (Chen et al., 2019b). Recently, CLPB was also found to have negative correlation with the progression-free survival in castration-resistant prostate cancer (Pudova et al., 2020). It is currently not clear why CLPB has such a broad role in different aspects of mitochondrial function, and how the ANK and NBD domains work together to fulfil its essential disaggregase activity. Here, we present a structural and functional characterization of human CLPB. Unexpectedly, we found that CLPB assembles into a homo-tetradecamers in the absence of substrate. Upon substrate binding, the CLPB complex is converted into dodecamers consisting two conventional hexameric units. The NBD ring within a hexamer shares common structural features of typical AAA+ unfoldase/disaggreagase, with a spiral arrangement of pore-loops to interact with a fully threaded substrate. The N-terminal ANK domain is essential for the higher-order organization of CLPB, and contributes to the disggregase activity by directly interacting with various mitochondrial substrates. These results provide a framework for further dissection of the role of CLPB in regulating mitochondrial functions.

## Results

### Cryo-EM structures of CLPB in the apo-state and substrate-bound state

We cloned the coding sequence of *CLPB* from a home-made cDNA library of HEK-293T cells, and sequencing result indicated that it was the splicing isoform 2 (UNIPROT: Q9H078-2). Different expression constructs were tested. Firstly, CLPB without the mitochondrial targeting sequence (MTS) (Wortmann et al., 2015), named CLPB- N92 (Fig. 1A), was expressed and purified from *E. coli* cells. Consistent with a previous study (Mroz et al., 2020), CLPB-N92 formed large aggregates and was highly heterogenous in size as shown by gel filtration and negative-staining electron microscopy (nsEM) (Supplementary Fig. 1A and B). In addition to the signal peptide removal, CLPB was reported to be further processed by a mitochondrial protease PARL localized in the inner membrane, and the predicted cleavage site is between C126 and Y127 (Saita et al., 2017). Notably, a hydrophobic stretch within the region removed by PARL (Fig. 1A) was reported to inhibit the disaggregase activity of CLPB (Cupo and Shorter, 2020). Therefore, as a validation, we expressed full-length CLPB exogenously in HEK-293T cells, and examined the sizes of protein products. Western blotting analysis showed two clear bands with the precursor form gradually decreasing over time (Supplementary Fig. 1C). Next, another construct harboring the sequence of CLPB starting from PARL cleavage site (named CLPB-N127) was tested for HEK-293T expression, which resulted in a protein product in the same size as the mature form of CLPB (Supplementary Fig. 1D). Therefore, the CLPB-N127 construct was finally used for CLPB protein preparation from *E. coli* cells.

Interestingly, both gel filtration and nsEM showed that the purified CLPB complex is in a higher oligomeric state rather than a hexamer as expected from typical AAA+ unfoldases and disaggregases (Supplementary Fig. 2A and B). And this higher-order organization is irrelevant of exogenously supplemented AMPPNP (Supplementary Fig. 2A). Despite this unusual observation, the purified CLPB complex is competent in both ATPase and disaggregase activities (Supplementary Fig. 1E and F). AMPPNP-treated CLPB complexes were then subjected to cryo-EM analysis. The 2D classification showed a three-layered architecture and a heptameric feature for the side-view and top-view average images, respectively (Fig. 1B). Further 3D classification indicated that CLPB complexes are double-heptamers, and they are extremely dynamic in structure, with both inter- and intra-heptamer conformational variations. With different 3D classification strategies, we could only push the overall resolution to a range of 6-7 Å (Supplementary Fig. 3, S1 Table). At this resolution, secondary structures were resolved in certain regions, and the map matches well with the predicted model of CLPB by AlphaFold (Jumper et al., 2021). The overall size of the complex is 175 Å and 160 Å in height and width, respectively. The seven NBDs form a closed ring, with an open channel (24 Å in diameter) in the center. The “head-to-head” organization of the two heptamers is mediated by their N-terminal ANK domains (Fig. 1C). Although a heptameric form of the ATPase modules is rare among AAA+ ATPases, another mitochondrial AAA+ ATPase Bcs1 was recently reported to exist in a homo- heptamer form (Kater et al., 2020; Tang et al., 2020).

Since the double-heptamer contains no substrate and is widely open in the central pore, we set out to obtain a structure of the substrate-engaged CLPB complex. A convenient way of doing this is through a Walker B mutation (E425Q) in the NBD domain. For most of the AAA+ proteins, this mutation would greatly slow down ATP hydrolysis but not ATP binding. Structural studies of a few AAA+ unfoldases/disaggregases showed that this mutation often resulted in a co-purification of endogenous peptide in the central channel (Fei et al., 2020; Lo et al., 2019; Twomey et al., 2019). Therefore, we prepared CLPB^E425Q^ mutant from *E. coli* cells (Supplementary Fig. 2C and D) and analyzed the sample by cryo-EM (Supplementary Fig. 4). Unexpectedly, top-view class averages from 2D classification showed a hexameric ring (Fig. 1B and Supplementary Fig. 5B). Similar to the wild type (WT) CLPB complex, the mutant complex is also highly dynamic and the global density map could only be refined to a global resolution of 7.9 Å. With local refinement, the NBD ring was improved to 5.2 Å (Supplementary Fig. 4C, D and F). From the structure, it is clear that the mutant CLPB^E425Q^ now takes a double-hexamer (dodecamer) form. Compared with the WT structure, while the width of the mutant complex reduces from 160 Å to 135 Å, the overall height increases from 175 Å to 200 Å (Fig. 1D). This is due the spiral configuration of CLPB^E425Q^ subunits within each hexamer. From the improved map of the NBD ring, the substrate density in the central channel could be unambiguously identified (Supplementary Fig. 4D). In general, the helical arrangement of subunits within a hexamer is highly similar to other substrate-bound AAA+ ATPases (Cooney et al., 2019; Gates et al., 2017; Twomey et al., 2019).

To confirm that this higher-order organization of CLPB is not an artifact of exogenous expression in *E. coli*, we examined the oligomeric states of CLPB variants purified from HEK-293T cells. The nsEM analysis indicates that both the WT and mutant CLPB complexes derived from human cells display a similar higher-order organization as the *E. coli* ones (Fig. 2A). Next, to rule out the possible splicing isoform specific effect on the oligomeric state, we also examined the isoform-1 of CLPB (UNIPROT: Q9H078-1) purified from *E. coli* cells (Supplementary Fig. 2E-F) using both nsEM and cryo-EM. This isoform-1 is 30 residues longer in the ANK domain than the isoform-2. Our results show that CLPB^isoform1^ complexes are again double-heptamers with an open central channel in the NBD ring (Fig. 2E, Supplementary Fig. 5C).

**Figure 2.**
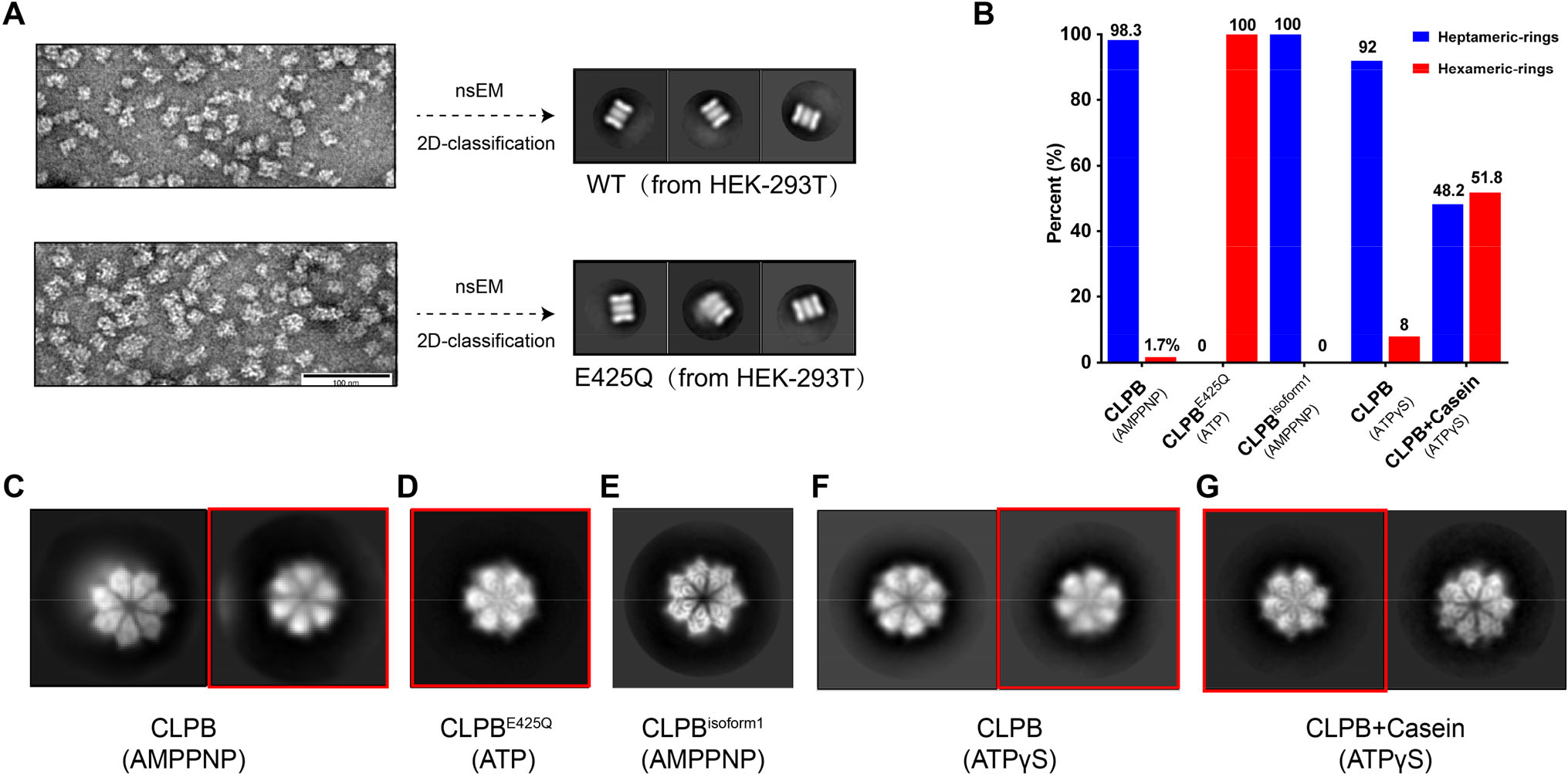
The CLPB double-heptamers transform into double-hexamers upon substrate binding. **A** CLPB complexes purified from HEK293 cells form double oligomeric state. Representative nsEM images (left) and 2D classification averages of nsEM particles (right) of CLPB and CLPB^E425Q^ from HEK-293T cells. **B** The proportion of the hexameric and heptameric top views in the CLPB-AMPPNP, CLPB^E425Q^-ATP, CLPB^isoform1^-AMPPNP, CLPB-ATPγs and CLPB+Casein-ATPγs datasets. **C-G** Representative top views of 2D classification averages of CLPB-AMPPNP (C), CLPB^E425Q^-ATP (D), CLPB^isoform1^-AMPPNP (E), CLPB-ATPγs (F) and CLPB+Casein-ATPγs (G) datasets. The top views with hexameric feature are indicated by red boxes.

In short, these results demonstrate mitochondrial CLPB, unlike other typical AAA+ proteins, adopts a unique higher-order structure. The CLPB-specific ANK domain mediates the attachment of two heptamers/hexamers through a head-to-head dimerization. Both the double-heptamer and double-hexamer are highly dynamic. Several factors contribute to the conformational flexibility of the full-length complexes. The first is the hexamer/heptamer interface. Although the ANK domains from the two heptamer/hexamers interact with each other, they do not have lateral interactions. Thus, the interface between two heptamers/hexamers is rather flexible. The second is the flexibility within each heptamers/hexamers, because the AAA+ domain rings are known to adopt unsymmetrical and dynamic conformations.

### Substrate binding induces the transition from double-heptamer to double-hexamer

A major difference between the WT and E425Q CLPB structures is the presence of a peptide substrate in the central pore, which brings close CLPB subunits to form a more compact structure. As shown in the 2D classification results, most of the top-view classes of the WT CLPB particles are heptameric, and only a tiny top-view class, 1.7% of all top-view particles, display a hexameric feature (Fig. 2B-C, Supplementary Fig. 5A). In contrast, all the top-view classes of the CLPB^E425Q^ particles are hexameric exclusively (Fig. 2B-D, Supplementary Fig. 5B). This suggests that the binding of substrate may induce a transition from tetradecamers to dodecamers.

To test this hypothesis, we incubated the WT CLPB complexes with a model substrate casein (Cupo and Shorter, 2020) in the presence of excessive ATPγS, which has been shown to best promote the binding of casein to Hsp104 (Gates et al., 2017; Weaver et al., 2017). As a control, WT CLPB complexes were also treated with ATPγS alone. Cryo-EM 2D classification was employed to analyze their oligomeric states. For the CLPB-ATPγS dataset, only a small fraction (8.0% of all top-view particles) shows a hexameric ring (Fig. 2B and F), indicating that the majority of particles retain the form of double-heptamer. In sharp contrast, in the presence of casein, the percentage of hexameric top-view particles has increased to 51.8% (Fig. 2B and G). These results indicate that substrate binding is likely the most important factor that drives the formation of double-hexamers from double-heptamer.

### The ANK domain is essential for the assembly and function of CLPB complexes

To validate the role of the ANK domain in organizing the higher-order structure, we created an NBD-only variant by truncating the N-terminal ANK domain. Gel filtration and nsEM reveal that CLPB-NBD proteins still form oligomers (Supplementary Fig. 9), but the size is much smaller than the double-heptamer or double-hexamer. Thus, these results experimentally proved a role of the ANK domain in assembling the higher-order CLPB complex.

Next, we examined whether these CLPB-NBD oligomers have ATPase and disaggregase activities. Our results show that the disaggregase activity critically depends on the integrity of both the ANK and NBD domains. The introduction of E425Q mutation (CLPB^E425Q^ or CLPB-NBD^E425Q^) and the deletion of ANK domain (CLPB-NBD) both resulted in a complete loss of the disaggregase activity. This is consistent with a previous study (Cupo and Shorter, 2020), suggesting an essential role of the ANK domain in the disaggregase function of CLPB. However, in contrast to the same study, our data show that CLPB-NBD is active in hydrolyzing ATP, and the activity is even higher than the WT CLPB (Supplementary Fig. 6). This discrepancy is due to the fact that CLPB-NBD in that study only formed dimer-like species under their pH 8.0 condition, whereas we used an acidic condition (pH 6.8) that is more relevant to that of the mitochondrial IMS.

For functional analysis of the ANK domain, we determined the crystal structure of the ANK domain at 2.1 Å resolution (S2 Table). The ANK domain of CLPB contains four ankyrin motifs (AM) and a unique insertion (consisting of two short helices) between AM2 and AM3 (Fig. 3A, right panel). The isoform 1 of CLPB differs from the isoform 2 exactly in this region, with 30 residues more in this insertion sequence (Fig. 3A, left panel). Since the ANK domain is important for both the assembly and function of CLPB, we wonder whether this insertion is functionally relevant. From the density map of the double-hexamer, the substrate density extends from the RBD ring to the layers of ANK domains, and some extra density in the center of the ANK layers was observed (Supplementary Fig. 7A). Fitting of the crystal structure of the ANK domain reveals that the insertion exactly locates in the inner surface of the ANK layers. Interestingly, the insertion contains two patches of hydrophobic residues (Fig. 3B), implying a potential role in substrate binding. Therefore, we constructed another CLPB mutant with the insertion completely deleted (CLPB^Δ201-232^), and compared this mutant with both isoforms 1 and 2 in their ATPase and disaggregase activities. The CLPB^Δ201-232^ mutant showed the same oligomeric state as the isoforms 1 and 2 (Supplementary Fig. 7C to E). The two isoforms have comparable activities in these two molecular functions (Fig. 3C). Although the deletion in CLPB^Δ201-232^ had no effect on the ATPase activity, it reduced disaggregase activity sharply by more than 50% (Fig. 3C and D). Therefore, this insertion is likely involved in the substrate recognition and may also contribute to the translocation of the substrate to the C-terminal NBD ring.

**Figure 3.**
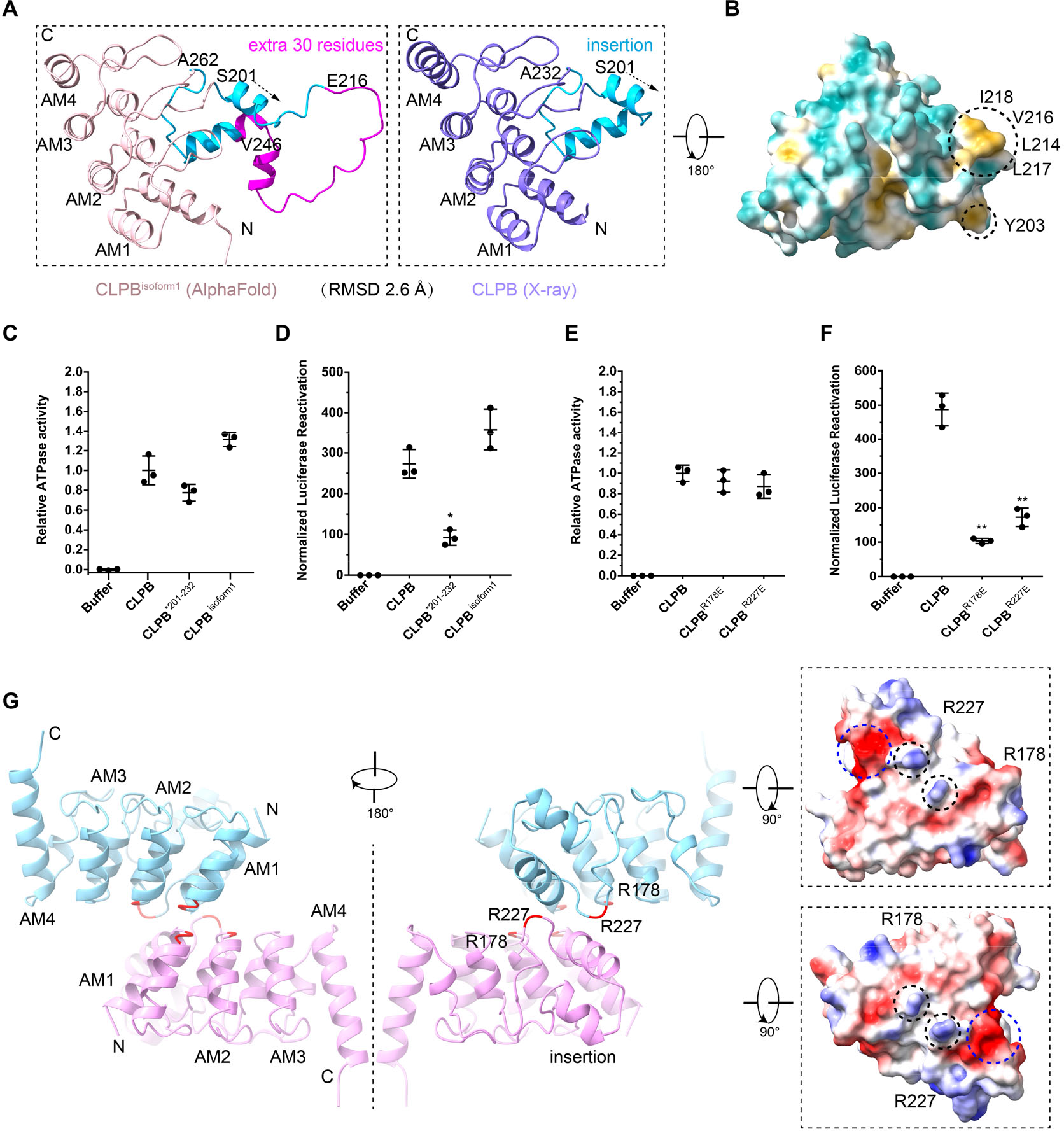
ANK domain is essential for the disaggregase activity of CLPB. **A** X-ray crystallography structure and AlphaFold predicted model of the ANK domain. The RMSD between these two structures is 2.6 Å. **B** The hydrophobic surfaces of the ANK domain. Two hydrophobic patches were found in the unique insertion. **C** The ATPase assay of CLPB, CLPB^Δ201-232^ and CLPB^isoform1^. ATPase activity was compared to CLPB (N=3, individual data points shown as dots, bars show mean ± SD). **D** The disaggregase activity assay of CLPB, CLPB^Δ201-232^ and CLPB^isoform1^. Disaggregase activity was compared to CLPB (N=3, individual data points shown as dots, bars show mean ± SD, *p<0.05). **E** ATPase assay of CLPB, CLPB^R178E^ and CLPB^R227E^. **F** Disaggregase activity assay of CLPB, CLPB^R178E^ and CLPB^R227E^. Results show that the disaggregase activities of CLPB^R178E^ and CLPB^R227E^ are reduced by 70-80% (N=3, individual data points shown as dots, bars show mean ± SD, **p<0.01). **G** The ANK domain dimer interface. One is the AM1-AM2 and the other is the second helix of the insertion. Two positive residues (R178 and R227) are outstanding in the ANK surface electrostatic potential.

Next, we analyzed the dimer interface of the ANK domain by docking the ANK models into the density maps of the double-heptamer and double-hexamer (Supplementary Fig. 8A and B). The ANK dimers in the two higher-order structures are not identical but generally similar. For the ANK dimer in the double-hexamer, two regions of the ANK domain contribute to the dimerization, the AM1/2 motifs and the insertion (Fig. 3G). One interface is relatively extensive and formed by four connecting loops between the first and second helices of the AM1 or AM2 from the two opposite ANK domains. The other interface is mediated by the second helix (and its downstream flanking sequence) of the insertion, and two arginine residues, R178 and R227, appear to be important in this interface. Based on the electrostatic surface potential, these two residues of one ANK domain point to a highly negatively charged surface patch (E140, E221, D222 and D223) of the opposite ANK domain (Fig. 3G). In fact, both R178 and R227 are highly conserved among the metazoan species (Cupo and Shorter, 2020). Therefore, we performed mutagenesis (R178E and R227E) to test whether they could disrupt the ANK dimerization. Unexpectedly, the two CLPB mutants still maintain a higher-order assembly, but their disaggregase activities are severely impaired by 70-80% (Fig. 3E and F, and Supplementary Fig. 8C-E). These results further emphasize a role of the ANK domain in the disaggregase function of CLPB.

Altogether, we demonstrate that the ANK domain is essential for both the assembly and function of the CLPB complex, and the functional importance of the ANK domain could be largely attributed to its insertion sequence between the AM2 and AM3.

### CLPB-NBD assembles into polypeptide-engaged helical structures

Due to the large inter- and intra-hexamer/heptamer flexibility, the structures of full-length CLPB complexes were not solved in atomic resolution. CLPB-NBD was thus used as a surrogate for high-resolution structural determination. Two mutant versions of CLPB (NBD and NBD^E425Q^) were analyzed by cryo-EM (Supplementary Fig. 9A-F), which again confirmed that E425Q mutation resulted in co-purification of an endogenous peptide in the central channel. Also similar to the full-length complexes, the top-view classes of the NBD^E425Q^ particles are exclusively hexameric, whereas both hexameric and heptameric arrangements were observed for the NBD^WT^ particles (Supplementary Fig. 9F). Subsequently, we focused on the NBD^E425Q^ dataset for high-resolution refinement. After several rounds of 3D classification, three different oligomeric arrangements, hexameric, heptameric and nonameric, were identified. The nonamer appeared to be more stable and could be resolved at an overall resolution of 3.7 Å (Supplementary Fig. 10), allowing the atomic modelling of the NBD of CLPB. In the nucleotide-binding pockets of the nonamer, a total of eight ATP molecules could be modelled (Supplementary Fig. 11). The conserved functional motifs on the ATPase domain are well resolved, including the Walker A motif (351-GSSGIG<u>K</u>T-358), sensor-1 (464-TS<u>N</u>-466), and sensor-2 (588-GA<u>R</u>-590) (Fig. 5E). The conserved residues I317, I318 and F541 form a hydrophobic pocket to stabilize the adenine ring of ATP, while T358 stabilizes the β- and γ-phosphates through a Mg^2+^ ion (Fig. 5E). The Arginine Finger (R531) from the adjacent protomer points to the γ-phosphate of ATP. In general, these structures show that the NBD of CLPB is a typical AAA+ ATPase module, underscoring a conserved mechanism of ATP-powered substrate unfolding and translocation (Gates and Martin, 2020).

The central channels of all three states are occupied by a peptide substrate, which forms successive interactions with the helically arranged CLPB protomers (Fig. 4A-C). In the nonamer, 9 protomers form a 1.5-turn helix around the central substrate. Besides the protomer number, the three oligomers also show obvious structural differences. While the twist angle between neighboring subunits is roughly 60° in all three forms, the axial rise within each oligomer is not constant and varies in the hexamer and heptamer largely. The axial displacements between P1 and P6 are roughly 25 Å, 30 Å and 35 Å for the hexamer, heptamer and nonamer, respectively (Fig. 4D-F). This is expected, as more axial space is required to fit in the seventh and more subunits onto the hexamer. In the full-length complexes, the ANK domain provides a steric hindrance to end the helical extension.

**Figure 4.**
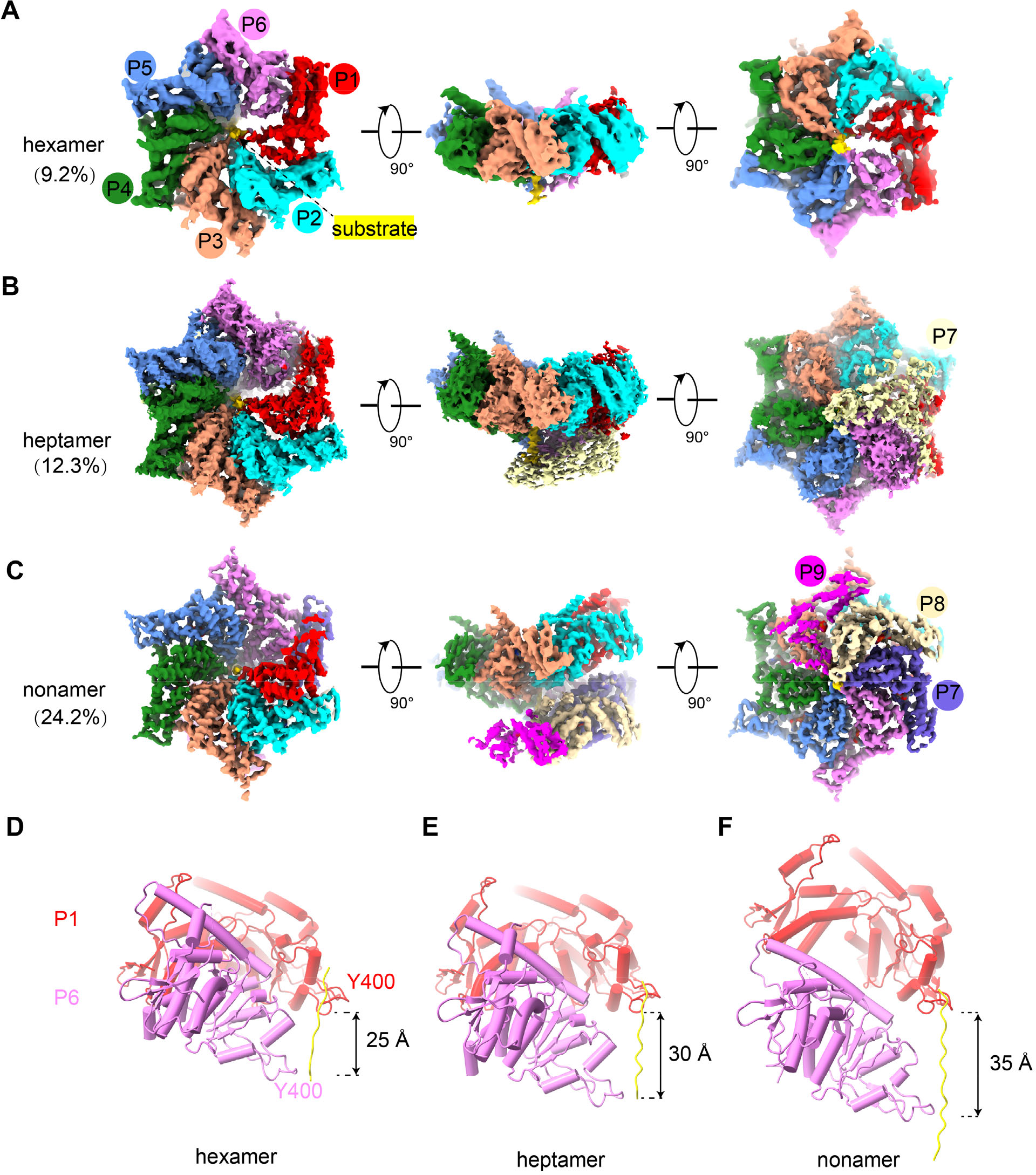
Cryo-EM characterization of the NBD helical structures of CLPB. **A-C** Density maps of the NBD hexamer (A), heptamer (B) and nonamer (C), respectively. Protomers are painted in different colors. **D-F** Distances along the central substrate between the pore-loop residue Y400 of P1 and P6 in the hexamer (D), heptamer (E) and nonamer (F). The axial rise of P6 relatively to P1 are labelled.

Overall, these data show that the ANK domain is essential in determining the oligomeric state of CLPB complex, and also explain why our CLPB-NBD still retains ATPase activity. Although this nonameric helical structure does not exist in a physiological context, it demonstrates that CLPB-NBD shares many characteristics of typical AAA+ proteins.

### The substrate interacts with conserved pore-loops of the helically arranged NBDs

The density of the polypeptide backbone is well resolved in the nonamer and was modeled as a 17-residue poly-alanine fragment (Fig. 5A and B). The peptide spans about 53 Å length in the central channel of the nonamer and displays successive interactions with the PL-I and PL-II from NBD protomers (Fig. 5B). For each protomer, a canonical PL-I (399-GYVG-402) binds to the peptide through two hydrogen bonds formed between the main-chain atoms. The first is between the carbonyl oxygen of the substrate residue N and the main-chain nitrogen of V401, and the other between the carbonyl oxygen of G399 and the main-chain nitrogen of the substrate residue N+2 (Fig. 5C). This pattern of interactions is nearly identical for all protomers, except that the hydrogen bond distances vary between 2.5-4.0 Å. Thus, the axial step size of PLs is exactly two residues. Of note, the main-chain atoms of Y400 do not participate in sequence-independent interaction with the substrate backbone. Therefore, the essential role of Y400 in the disaggregase activity should arise from its aromatic side-chain, which could interact with various side- chains of a threaded substrate. Moreover, the substrate could be modeled in both directions (N to C or C to N) in the density map, and both configurations could satisfy this repeated pattern of interactions (Fig. 5C and D). In contrast to the PL-I, the PL-II, consisting of E386, R387, and H388, is relatively away from the substrate backbone, except that the side-chain of H388 is within 4-Å distance with the β-carbon atom of the substrate (Fig. 5B). Thus, it is likely that the primary role of PL-II during substrate processing is to interact with different side-chains of the substrate through its polar residues.

**Figure 5.**
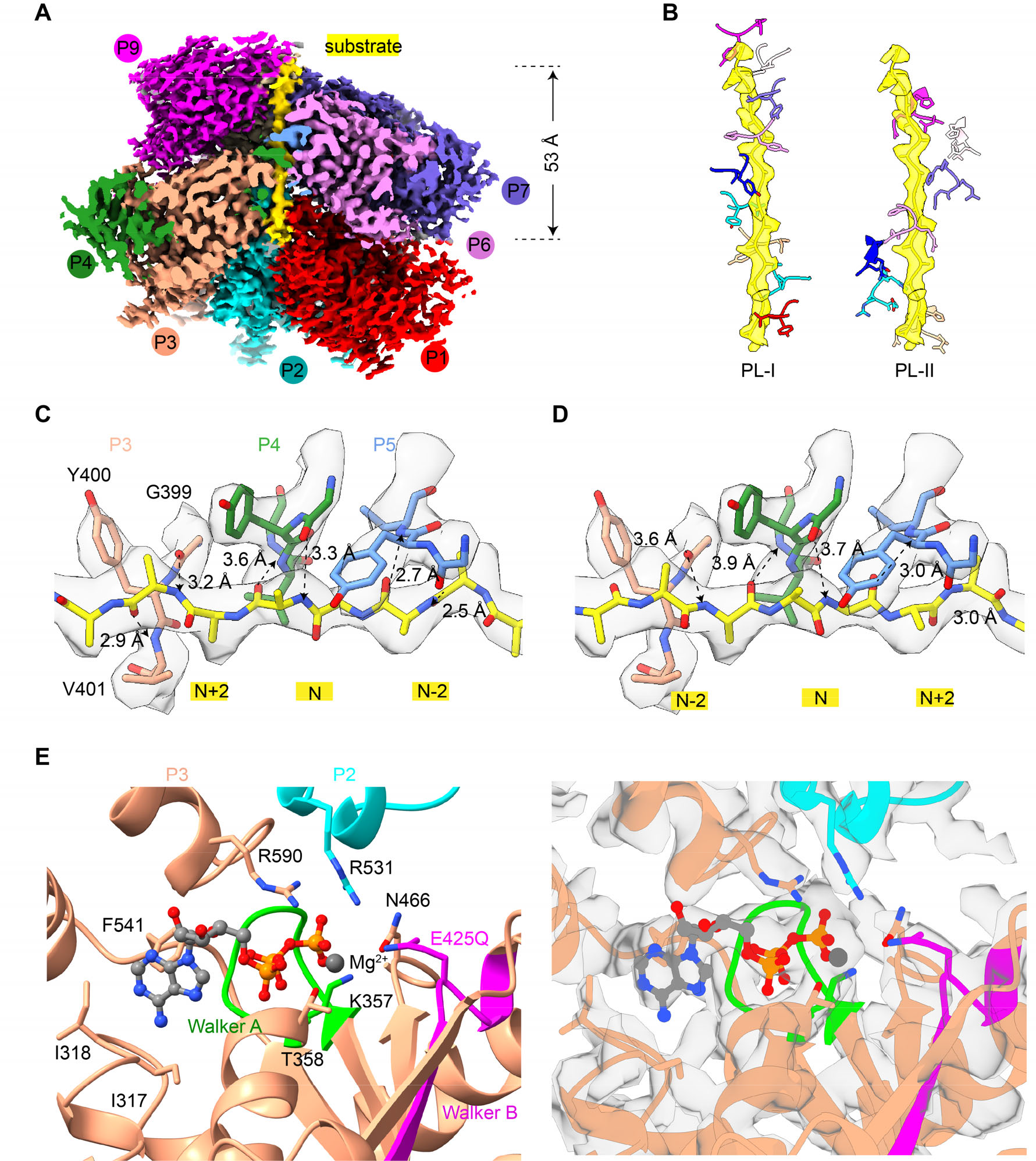
Structure of the NBD nonamer in processing a peptide substrate. **A** Density map of the NBD nonamer in the substrate-processing state. Nine protomers are indicated as P1 to P9 and painted in different colors. The central substrate is colored yellow. **B** The spiral configuration of the PLs of CLPB protomers around the substrate in the central channel. PL-Is and PL-IIs are shown in the left and right panels, respectively. While the conserved PL-Is (G399-Y400-V401-G402) directly interact with the substrate, the PL-IIs (E386-R387-H388) situate in slightly larger distances from the substrate. **C, D** Magnified view of the interactions between the PL-I of P3, P4 and P5 and the backbone of substrate. The substrate could be modeled in both directions, N to C or C to N. The potential hydrogen bonds are indicated by dashed lines and the distances are labelled. **E** Magnified view of the conserved ATP-binding pocket of CLPB. Functionally important residues of Walker A motif (K357, T358), Walker B motif (E425), sensor-1 (N466), sensor-2 (R590), arginine finger (R531) and the conserved residue I317, I318. Atomic model and density map are shown in the left and right panels, respectively.

In short, these results indicate that the NBD ring of CLPB is a typical unfoldase core, and the PLs of CLPB functions to grip and move the substrate within the central channel in a conserved manner as classic AAA+ unfoldases/disaggregases.

### Mitochondrial interactome analysis of the ANK domain

Previous data showed that *CLPB* knockout cells exhibited decreased solubility for many proteins in the IM and IMS of mitochondria, including HAX1, TOMM22/70, TIMM22/23, HTRA2, PHB1/2, OPA1, STOML2 and SLC25 family proteins (Cupo and Shorter, 2020). Given the essentiality of the ANK domain in the disaggregase activity of CLPB, we performed a mass spectrometry (MS) based interactome analysis of the ANK domain. A plasmid harboring the MTS and ANK domain sequences was expressed in HEK-293T cells, and mitochondria were then isolated, lysed and dissolved in 1% detergent. The ANK domain and its binders were purified through a C-terminal Strep tag, and subject to MS analysis. A relatively stringent criterion (fold-change > 4 and p-value < 0.01) were used for enrichment analysis (three biological replicates). With DAVID Bioinformatics Resources (Sherman et al., 2022), we restricted our analysis on mitochondrial proteins detected in the samples. Compared with the control sample, 934 mitochondrial proteins were significantly enriched in the ANK sample, and most of them are located in the IM and matrix (60.3%). Although CLPB is located in the IMS, it may also function to promote the solubility of certain membrane proteins with IMS-exposed domains. Therefore, we focused on proteins from the OM, IMS and IM. As a result, 381 such proteins were selected for further analysis, including all of those potential substrates previously reported to be most affected by *CLPB*-knockout (Cupo and Shorter, 2020). If these proteins were ranked by their abundances, three IMS proteins stood out in the top-20 list, including HAX1, OPA1 and AGK (Fig. 6C). Consistently, both HAX1 and OPA1 have been experimentally validated to interact with CLPB (Chen et al., 2019b; Fan et al., 2022). In general, the ANK interactome identified in this study agrees well with previous proteomic data based on full-length CLPB (Chen et al., 2019b; Fan et al., 2022; Wakula et al., 2020), indicating that the ANK domain has a direct role in substrate binding.

**Figure 6.**
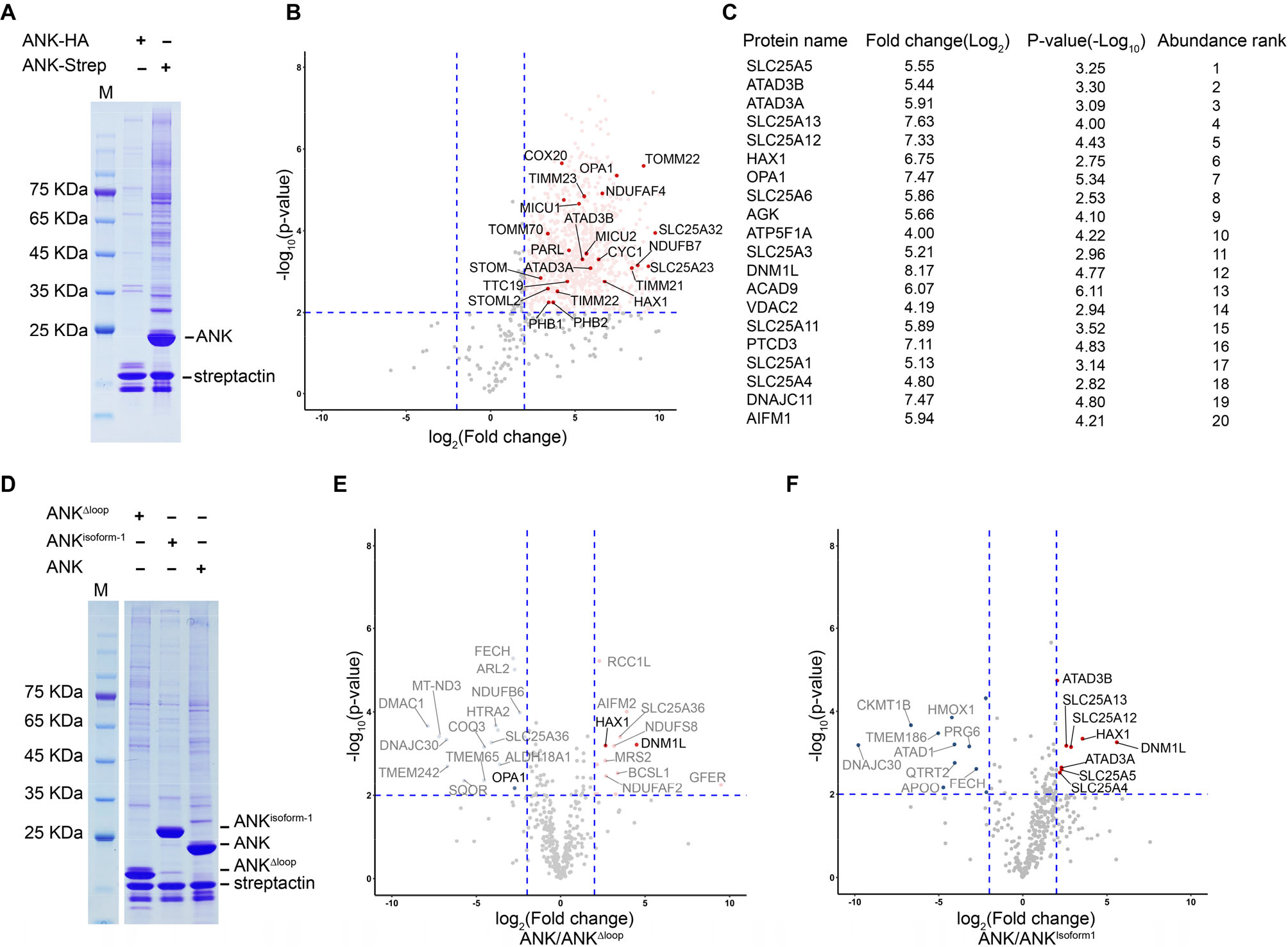
The mitochondrial interactome of the ANK domain. **A** HEK-293F cells expressing the MTS-ANK-HA or MTS-ANK-Strep constructs were subjected to anti-Strep immunoprecipitation. Precipitates were analyzed by SDS-PAGE, Coomassie blue staining and mass-spectrometry (MS). **B** Volcano plot showing the mitochondrial proteins co-precipitated with MTS-ANK-Strep. 934 proteins that were more enriched in the MTS-ANK-Strep are labelled in light red. The potential substrates, previously reported to be most affected by *CLPB*-knockout, are highlighted in red. A relatively stringent criterion (fold-change > 4 and p- value < 0.01) were used for enrichment analysis (three biological replicates), indicating with blue dashed lines. **C** Top-20 proteins in OM, IMS and IM are listed, based on the ranking of protein abundances. **D** HEK-293F cells expressing the MTS-ANK-Strep, MTS-ANK^Δloop^-Strep and MTS-ANK^isoform1^-Strep constructs were subjected to anti-Strep immunoprecipitation. Precipitates were analyzed by SDS-PAGE, Coomassie blue staining and MS. **E, F** Volcano plot showing the fold change of the OM, IMS and IM mitochondrial proteins in MTS-ANK-Strep compared to MTS-ANK^Δloop^-Strep (E) or MTS-ANK^isoform1^-Strep (F). Proteins in the top-20 list (C) were highlighted in red.

We also showed that the unique insertion between AM2 and AM3 in the ANK domain is important in the disaggregase activity *in vitro* (Fig. 3). Therefore, two ANK variants were next used for interactome analysis. The MTS-ANK^Δ201-232^ and MTS-ANK^isoform1^ were expressed in HEK-293T cells and their interactome were similarly analyzed by MS (Fig. 6D). Among the analyzed mitochondrial OM, IMS and IM proteins, eight proteins were more enriched in the ANK^isoform2^ sample, including ATAD3B/ATAD3A, SLC25A4/5/12/13, HAX1 and DNM1L. All of them are indeed among the most abundant top-20 list. In contrast, a larger number of proteins (fourteen) were selectively enriched in the ANK^isoform2^ sample than the ANK^Δ201-232^ sample, again including HAX1 and DNM1L (Fig. 6E, S6 Table). Very interestingly, a recent study reported that HAX1 preferentially binds to the isoform 2 of CLPB *in vivo*, rather than the isoform 1 (Fan et al., 2022), highly consistent with our findings.

These interactome analyses support a direct role of the ANK domain in substrate binding, and together with previous studies (Chen et al., 2019b; Cupo and Shorter, 2020; Fan et al., 2022) confirmed that HAX1 is one of the natural substrates of CLPB. The differential enrichment between the isoform1 and 2 further suggests a possible role of the ANK insertion in substrate selection, probably through altered binding affinity.

## Discussion

In the present work, we characterized the structures of CLPB in both the apo and substrate-bound states. Unexpectedly, we found that apo CLPB assembles into higher-order structures, in the form of a double-heptamer through inter-molecular interactions mediated by N-terminal ANK domains (Fig. 1). We further demonstrate that the double-heptamers could be efficiently converted to double-hexamers upon the addition of a model substrate (Fig. 2), highlighting the possibility that the double-heptamer is a physiologically relevant, resting state of CLPB in mitochondria. We believe that the heptameric form of CLPB is not a pure *in vitro* artifact for several reasons.

First, the heptamer form is the predominant species of apo CLPB (Fig. 2), indicating an intrinsic property of the CLPB ATPase module. Although AAA+ proteins are generally considered to be hexamers, an increasing number of AAA+ proteins were recently discovered to take heptamer as a primary oligomeric state (Hoffmann et al., 2022; Kater et al., 2020; Kim et al., 2020; Tang et al., 2020), including a mitochondrial IM-bound AAA+ protein Bcs1 (Kater et al., 2020; Tang et al., 2020). Second, The substrate-induced change on the oligomeric state has also been observed for other AAA+ members, such as DNA helicase AAV2 Rep68 (Santosh et al., 2020), RuvB (Miyata et al., 2000), and archaeal MCM (Yu et al., 2002). Third, the ATPase activity of full-length CLPB (double-hexamer) is lower than CLPB-NBD alone (Supplementary Fig. 6). In fact, roughly two third of CLPB-NBD particles exhibit a hexameric arrangement. This indicates that the heptamer is not optimized for efficient ATP-hydrolysis. Thus, as a substrate-free state, the heptameric form could avoid necessary ATP consumption in the IMS. Notably, the buffer we used to prepare CLPB complexes was at pH 6.8, which closely resembles that of the IMS (Santo-Domingo and Demaurex, 2012).

Nevertheless, with these said, the substrate-engaged CLPB is still hexameric as shown in the structures of CLPB^E425Q^ and NBD. In the high-resolution structure of the NBD^E425Q^ nonamer, the PL-Is of the nine protomers form a spiral staircase around the substrate in a two-residue step-size (Fig. 5C-D). There PL-Is interact with the substrate in a sequence-independent manner, and capable of accommodating polypeptide in both the N-C and C-N directions, implying that it could potentially thread the substrate in both directions, similar as bacterial ClpX and ClpA (Olivares et al., 2017). These data indicate that the mechanisms of CLPB in substrate threading and translocation are highly similar to those cytosolic AAA+ unfoldases/disaggregase, such as Cdc48/p97 and Hsp104 (Cooney et al., 2019; Gates et al., 2017; Pan et al., 2021; Twomey et al., 2019).

In the double-hexamer, both hexamers are in active conformation with the substrate being threaded through the central pores of their ATPase rings. This raises an interesting question: Whether or not the two hexamers work in a synergistic manner, since a CLPB hexamer already contains all essential structural features of a unfoldase/disaggregase. A conventional model is that the two hexamers work independently, and upon engagement with a protein aggregate the two hexamers do not need to maintain stable association for all six pairs of ANK domains. However, as seen in the structure of the double-hexamer (Fig. 1D), it is also possible that the axial movement of a CLPB subunit in one hexamer during the ATP-hydrolysis cycle could potentially affect another CLPB subunit in the other hexamer through their tightly associated ANK domains. Unfortunately, our attempt to obtain ANK mutants to separate its roles in structural organization and disaggregase activity has failed (Fig. 3E and F). Nevertheless, this “head to head” organization reminds us of the recent findings on the double-hexamer form of p97 (Caffrey et al., 2021; Gao et al., 2022; Hoq et al., 2021; Yu et al., 2021), although p97 utilizes a “back to back” interface through two C-terminal ATPase rings. These two examples underline a possibility that AAA+ proteins could adopt different higher-order assemblies to serve different purposes.

We also determined the crystal structure of the ANK domain, and analyzed its functional significance in the disaggregase function. Our data showed that the ANK domain is indispensable for the disaggregase activity of CLPB, and a deletion of the unique insertion in the ANK domain resulted in more than 50% reduction on the disaggregase activity (Fig. 3D). Furthermore, two single mutations of the ANK domain in the ANK dimer interface, including 178E and R227E, both greatly impaired the disaggregase activity by 70-80% (Fig. 3F). R227 is exactly located to the insertion sequence. These observations suggest that this unique insertion is functionally important. Since this insertion is at the innermost position of the ANK ring and distant from the pore-loops of the ATPase modules, the ANK domain may function to recognize and pass the substrate to the distal ATPase ring. This hypothesis was further supported by our interactome analysis of the separate ANK domain. A large number of the IM and IMS proteins were greatly enriched in the sample affinity-purified through tagged ANK, including all of those previously found to be prone to aggregation in Δ*CLPB* cells (Cupo and Shorter, 2020). Many of these candidates have also been reported to interact with CLPB in previous large-scale proteomics studies (Botham et al., 2019; Huttlin et al., 2021), and two of them have been experimentally validated, such as HAX1 and OPA1 (Chen et al., 2019b; Fan et al., 2022). Of note, several mitochondrial proteins involved in cristae remodeling are among the most enriched group, including OPA1, DNM1L, and IMMT. This appears to collaborate with the previous finding that CLPB loss resulted in altered cristae structure in mitochondria (Chen et al., 2019b). We also tested the ANK^Δ201-232^ and ANK^isoform1^ in interactome analysis. Compared with the ANK^isoform2^, a relative extensive change on the enrichment profile was observed for the ANK^Δ201-232^ sample. In contrast, comparison between isoform 1 and isoform 2 reveals that many potential substrates of CLPB are significantly more enriched in the sample of ANK^isoform2^, including HAX1 and DNM1L (Fig. 6E and F). These results further pinpoint a role of the ANK- insertion in the substrate recruitment, and suggest another level of regulation on CLPB function through alternative splicing.

The structures of CLPB also allow us to map disease related mutations (Capo-Chichi et al., 2015; Cupo and Shorter, 2020; Kanabus et al., 2015; Kiykim et al., 2016; Pronicka et al., 2017; Saunders et al., 2015; Wortmann et al., 2021; Wortmann et al., 2015) on the atomic model to understand their possible effect on the function of CLPB. Mutations associated with 3-MGA could be categorized into three classes. The first class of mutations lie at the interface between two adjacent CLPB protomers, including R378G, R445Q, Y587C, R598C and E609K (Fig. 7A). These mutations likely perturb the inter-protomer communication to impair CLPB function. The second class consists of M381I, C456R, E471K, and Y537C in the large subdomain, and A561V, G616V, R620P, and I652N in the small subdomain (Fig. 7B). These mutations are away from the inter-protomer or inter-domain interfaces, and they may destabilize the domain structure of CLPB (Fig. 7B). The third class of mutations are in the AM3 of the the ANK domain. Besides two nonsense mutations (R250* and K321*), three mutations (T238M, A239T and Y242C) are on the first helix of the AM3 (Fig. 7C). This region contains a characteristic tetrapeptide (-TPLH-) motif (238-TALH-241 in CLPB), and is highly conserved among ankyrin motifs (Li et al., 2006). Therefore, these mutations might destabilize the structure of the ANK domain, or lead to misfolding of the ankyrin motifs. Consistently, it was shown that T238M did not affect the ATPase activity, but largely inhibited the disaggregase activity (Cupo and Shorter, 2020). As for mutations related to SCN, they are exclusively located close to the active center of the NBD (Fig. 7D). They are distributed in sensor-1 (N466K), sensor-2 (R590C), AF (R531Q/G), and Walker A motif (T358K). It is apparent that they may affect the binding and hydrolysis of ATP (Fig. 5E). Indeed, CLPB with these mutations were defective in both the ATPase and disaggregase activities *in vitro* (Warren et al., 2022).

**Figure 7.**
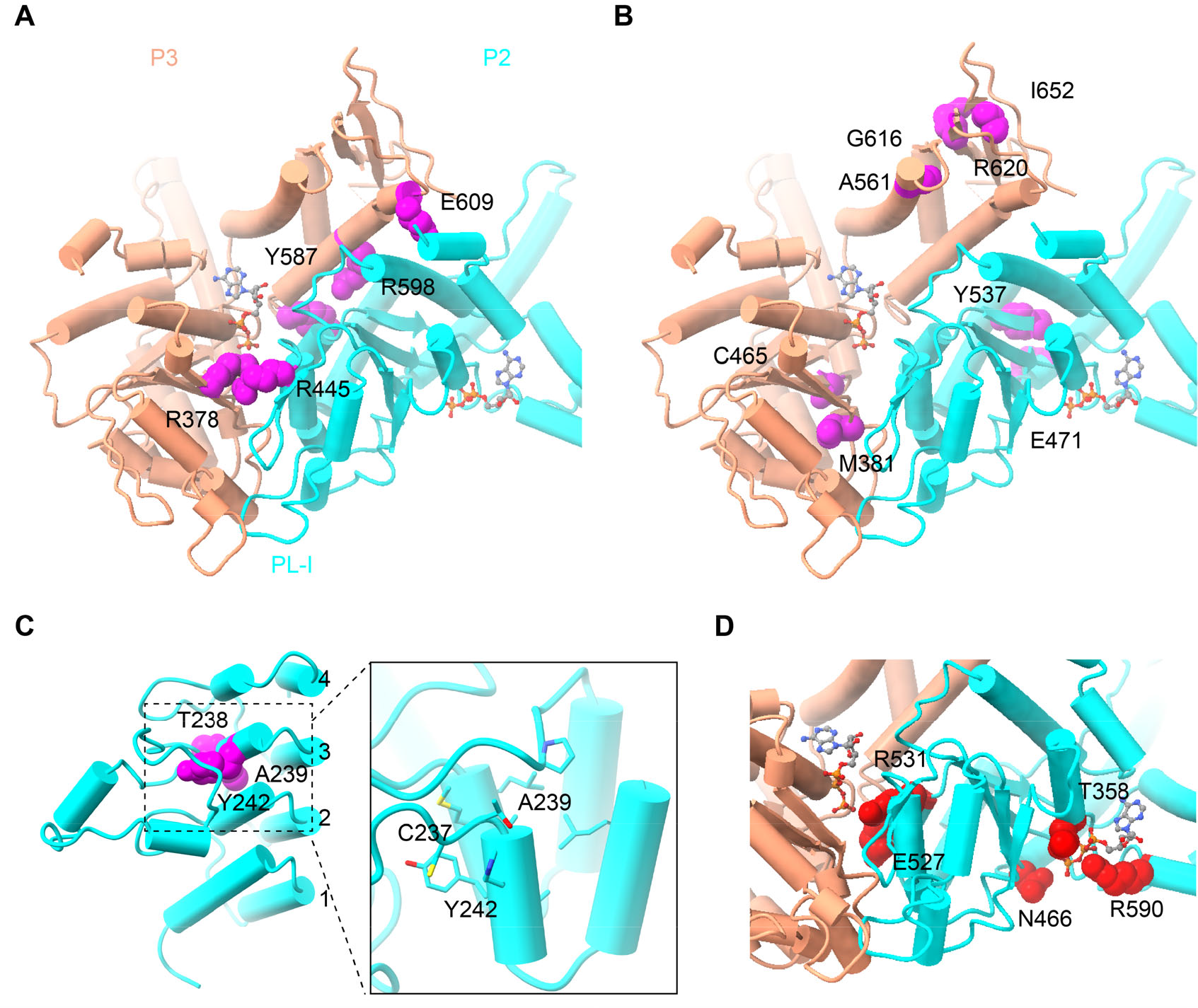
The disease-related mutations of CLPB in 3-MGA and SCN. **A** 3-MGA-related mutations in the interface of adjacent protomers. **B** 3-MGA-related mutations within the large and small subdomain of the ATPase domain. **C** 3-MGA-related mutations in the ANK domain. **D** SCN-related mutations in the ATP-binding pocket. Residues on the positions of these mutation are highlighted in magenta or red sphere models. ATP molecule is highlighted in stick models.

## Methods

### Protein expression and purification

The coding sequence (isoform 2) of human *CLPB*, amplified from a home-made cDNA library of HEK-293T cells, was cloned into pET28a vector with a N-terminal His6-SUMO tag followed by a TEV protease cleavage sequence, and expressed in *E. coli* Transetta (DE3) cells (Trans). CLPB protein variants were purified as reported previously (Mroz et al., 2020), with some modifications. Briefly, the culture was induced with 1mM IPTG at OD_600_ of 0.6-0.8 at 20°C overnight. Cells were harvested and resuspended in lysis buffer (50 mM Tris-HCl, pH 8.0, 500 mM NaCl, 5% glycerol, 1% Triton X-100 and 1mM PMSF) and lysed by sonication. Lysates were cleared by centrifugation in a JA-25.50 rotor (Beckman) for 30 min at 20,000 r.p.m. and the supernatants were precipitated with ammonium sulfate at 35% saturation, then incubated for 10 min at room temperature (RT). The cloudy lysates were centrifuged for 30 min at 15,000 r.p.m., and then dissolved the protein pellet with Buffer B (50 mM Tris-HCl, pH 8.5) supplemented with 5 mM DTT and centrifugation again. The solutions were loaded onto a Mono Q column (GE Healthcare). Proteins were eluted with a linear gradient to 60% Buffer C (50 mM Tris-HCl, pH 8.5, 1M NaCl). To remove His6-SUMO tags, TEV protease was added into the eluates, and then incubated at RT for 2 hrs. The tag and contaminations were removed by size-exclusion chromatography, which was pre-equilibrated with 20 mM HEPES-KOH, pH 6.8, 300 mM KCl, 5 mM MgCl_2_. Peak fractions were pooled and concentrated, then incubated with 5 mM nucleotide (AMPPNP for CLPB, ATP for CLPB^E425Q^) at RT for 2 hrs. 1 mM BS3 was added to improve the sample stability, and loaded onto size-exclusion chromatography. Peak fractions were first examined by nsEM, then pooled and concentrated. 1mM nucleotide was added before vitrification. CLPB^E425Q^ and CLPB^Y400A^ were prepared by site-specific mutagenesis, and purified as described above.

The NBD and NBD^E425Q^ sequences were cloned by site-specific mutagenesis. Harvested cells were resuspended in lysis buffer II (50 mM Tris-HCl, pH 7.4, 300 mM NaCl, 5% glycerol, 1% Triton X-100, 1 mM PMSF, and 20 mM imidazole) and lysed by sonication. The supernatants were incubated with Ni–NTA agarose beads (GE Healthcare) at 4°C for 2 hrs. After washed 5 times with lysis buffer, protein was eluted with elution buffer (50 mM Tris-HCl, pH 7.4, 300 mM NaCl, 5 mM MgCl2, and 500 mM imidazole). TEV protease was add into the eluates to remove His6-SUMO tags, and then incubated at RT for 2 hrs. Then the proteins were loaded onto size-exclusion chromatography, which was pre-equilibrated with 20 mM HEPES-KOH, pH 6.8, 300 mM KCl, 5 mM MgCl_2_. Peak fractions were pooled and concentrated, then incubated with 5 mM nucleotide (AMPPNP for NBD, ATP for NBD^E425Q^) at RT for 2 hrs, and then loaded onto size-exclusion chromatography. Peak fractions were first examined by nsEM, then pooled and concentrated. 1mM nucleotide was added before vitrification.

For the ANK domain purification, cells were resuspended in lysis buffer III (50 mM Tris-HCl, pH 8.0, 500 mM NaCl, 5% glycerol, 1% Triton X-100, 1 mM PMSF, and 20 mM imidazole). The supernatants were incubated with Ni–NTA agarose beads (GE Healthcare) at 4°C for 2 hrs. The eluates were loaded onto size-exclusion chromatography, and eluted with 50 mM Tris-HCl, pH 8.0, 150 mM NaCl.

### ATPase activity measurement

0.1 μM CLPB and CLPB variants were incubated with 1 mM ATP at 30°C for 30 min in ATPase activity buffer (AAB: 20 mM HEPES-KOH, pH 6.8, 150 mM KAOc, 5 mM MgCl2, 0.1% Tween-20). ATPase activity was measured by using the Malachite green phosphate assay (Lanzetta et al., 1979). For each protein at least two biological replicates were measured with three independent replicates.

### Luciferase disaggregation assay

Disaggregase activity was measured as previously reported (Cupo and Shorter, 2020). Briefly, 1 μM CLPB and CLPB variants were incubated with 50 nM fresh firefly luciferase aggregates at 30°C for 90 min in luciferase reactivation buffer (LRB: 20 mM HEPES-KOH, pH 8.0, 150 mM KAOc, 10 mM KAOc, 10 mM DTT) supplemented with 5 mM ATP. Notably, the disaggregase activity of CLPB was detected using the luciferase reporter assay kit (Beyotime, CAT# RG005) alone (Supplementary Fig. 1F). Others were detected with the luciferase reporter assay kit (Trans, CAT# FR101-01).

### Transient overexpression of CLPB variants

HEK-293 T/F cells were transfected with pCAG vector containing CLPB variants. Cells were transfected using pEI transfection reagent. After 48 hrs, cells were collected.

### Western blots

Cells were washed with PBS and incubated with lysis buffer B (50 mM Tris-HCl pH 7.4, 150 mM NaCl, 1% Triton X-100, 0.1 % SDS and 1mM PMSF) for 30 min on ice. α-Strep antibody (Huaxing Bio, CAT# HX1816) was used to detect the target protein.

### Mitochondrial isolation

Mitochondria were isolated as previously described (Cupo and Shorter, 2020; Frezza et al., 2007). Briefly, 800 mL cells were resuspended in 40 mL mitochondria isolation buffer (20 mM HEPES-KOH, pH 7.6, 250 mM sucrose and 2 mM EDTA) and homogenized with a Dounce homogenizer at 4°C. Lysate was then centrifuged at 1,300g for 5 min. The supernatant was collected and then centrifuged at 13,000g for 15 min twice. And the pellet was resuspended with lysis buffer C (20 mM Tris-HCl pH 8.0, 150 mM NaCl, 1% Triton X-100, and 1mM PMSF) and incubated for 30 min at 4°C. Lysate was then centrifuged at 20,000g for 30 min. The supernatant was collected and incubated with Strep resin for 2 hrs at 4°C. After extensively washed, SDS-loading buffer was added to the sample. The samples were separated by MOPS-PAGE on a gradient gel (4%-20%).

### LC-MS/MS analysis

The samples were excised into several parts form protein gels, reduced with DTT, alkylated with iodoacetamide (IAA), and subsequently digested with trypsin. Peptides were analyzed by Thermo Orbitrap Exploris 480 mass spectrometer (Thermo Fisher Scientific) coupled with an Easy LC system (Thermo Fisher Scientific). Peptides mixtures were loaded onto C18 Trap Column in Buffer A (0.1% formic acid) and separated with a gradient Buffer B (0.1% formic acid, 80% ACN) (2 min 4-8% B; 37 min 8-25% B; 11 min 25-35% B;7 min 35-95% B; 3min 95% B) at a flowrate of 300 nL/min. Data were acquired in data-dependent mode with s with the following settings: MS1 60,000 resolution, 300 to 1650 m/z of mass range; MS2 30% normalized collision energy, 15,000 resolution, 1.6 m/z of isolation window. Proteins were identified and quantified by Proteome Discoverer software 2.2 using default settings against a database of UNIPROT *Homo sapiens*. Methionine oxidation and N-terminal acetylation were set as variable modifications. Protein level and peptide level false discovery rate (FDR) was set at 1%.

The differential enrichment analyses were based on three biological replicates (starting from expression plasmid transfection), and a stringent screening condition (p-value < 0.01 and Fold change > 4) was used. In the MTS-ANK- Strep samples, 1,115 mitochondrial proteins were detected, of which 934 proteins were highly enriched, compared with the MTS-HA sample (S3 Table). From these highly enriched proteins, those located in the OM, IMS and IM were selected for further analysis (S5 Table, Fig 6C). The comparison between the MTS-ANK-Strep and MTS- ANK^Δ201-232^-Strep samples was focused on 374 mitochondria proteins with OM, IMS and IM locations (S6 Table). The comparison between the MTS-ANK-Strep and MTS-ANK^isoform1^-Strep samples was similarly done, and 332 OM, IMS and IM proteins were used for analysis.

### Cryo-EM specimen preparation

The sample homogeneity was first examined by nsEM. Proteins were applied onto carbon-coated copper grids, and examined with an FEI Tecnai T20 TEM at 120 kV. During specimen preparation for cryo-EM, CLPB and CLPB^E425Q^ particles displayed strong preferred orientation. 0.5 mM CHAPSO was added to reduce the preferred orientation (Chen et al., 2019a). Cryo-grids (R1.2/1.3, Au300, Quantifoil) were prepared with an FEI Vitrobot Mark IV with the chamber set at 6°C and 100% humidity. Grids were screened with an FEI Talos Arctica before data collection.

### Cryo-EM data collection and image processing for CLPB

For the sample of CLPB-AMPPNP, micrographs were recorded with 300 kV FEI Titan Krios (Gatan K3 direct electron detector) with GIF Quantum energy filter (Gatan) at a nominal magnification of 81,000X and at a calibrated pixel size of 1.07 Å. 40 frames with exposure time of 3.2 s were collected with defocus ranged from -1 to -1.4 μm under low-dose condition. A total of 9,341 micrographs from two datasets were collected with EPU. The micrographs were subjected to beam-induced motion correction using MotionCor2 (Zheng et al., 2017), and contrast transfer function (CTF) parameters for each micrograph were determined by Gctf (Zhang, 2016). RELION (versions 3.1) was used to perform data processing (Zivanov et al., 2020). After auto-picking, the particles were subjected to 2D classification and several round 3D classifications. 299,306 particles from two datasets were selected and applied for the next round of 3D classification (K=4), a final set of 231,212 particles were kept for 3D refinement. For double-heptamer, another round of 3D classification (K=6) without alignment was performed to improve the resolution. Finally, 153,005 particles were kept for the final 3D refinement (final resolution 6.8 Å). Due to the inter-heptamer flexibility, mask-based (only including a single heptamer) classification (K=6) and refinement was applied, which showed no significant improvement (Supplementary Fig. 3). In addition, multiple rounds of focused classification based on the NBD-ring or a selected combination of CLPB subunits using symmetry expansion derived half-particles were also tried, with different setting of parameters, however, none of them was able to reach better than 5 Å resolution.

For the sample of CLPB^E425Q^-ATP, micrographs were recorded with 300 kV FEI Titan Krios (Gatan K2 camera) with GIF Quantum energy filter (Gatan) at a nominal magnification of 36,000X and at calibrated pixel sizes of 1.052 Å. 32 frames with exposure time of 8 s were collected with defocus ranged from -1.2 to -1.8 μm under low-dose condition. A total of 4,574 micrographs were collected with SerialEM (Mastronarde, 2005). After 2D classification and several round 3D classifications, a 7.5-Å density map was achieved (Supplementary Fig. 4). In Addition, a density map of 5.2-Å could be achieved by applying a local mask of the NBD-ring during reconstruction (Supplementary Fig. 4C, D and F).

For the sample of CLPB^isoform1^-AMPPNP, a total of 953 micrographs were recorded with 200 kV FEI Titan Krios (Gatan K2 camera) at a magnification of 36,000X and at calibrated pixel sizes of 1.157 Å. 32 frames with exposure time of 8 s were collected with defocus ranged from -1.1 to -1.8 μm under low-dose condition. After several rounds of 2D classifications, only hexameric rings of top views were observed (Supplementary Fig. 5C). For the sample of CLPB-casein-ATPγs, 20 μL casein (4.0 mg/mL) was added into 40 μL CLPB (2.0 mg/mL) before vitrification. A total of 885 micrographs were recorded with 200 kV FEI Titan Krios (Gatan K2 camera) at a magnification of 36,000X and at calibrated pixel sizes of 1.157 Å. 32 frames with exposure time of 8 s were collected with defocus ranged from -1.4 to -2.0 μm under low-dose condition. After several rounds of 2D classifications, 54,771 particles were left, and hexameric rings (25,867 particles) and heptameric rings (24,046 particles) of top views were both observed (Supplementary Fig. 5D). As a control, CLPB-ATPγs dataset was prepared as CLPB-casein-ATPγs without adding casein.

For the sample of NBD^E425Q^-ATP, micrographs were recorded with 300 kV FEI Titan Krios G3i (Gatan K3 direct electron detector) with GIF Quantum energy filter (Gatan) at a magnification of 64,000X and at pixel size of 1.08 Å. 32 frames with exposure time of 2.44 s were collected with defocus ranged from -1 to -1.4 μm. A total of 4,574 micrographs were collected. 3,753,541 particles were split into 2 parts to facilitate processing. After several rounds of 2D classification and 3D classification, 3 major classes were found: hexamer, heptamer, and nonamer. A small group of hexamer particles were used for 3D refinement, resulting in a density map of 7.4 Å (Supplementary Fig. 10). The particles of heptamer and nonamer were subjected additional rounds of 3D classification, and after CTF refinement and Bayesian polishing, were reconstructed to the final resolutions of 4.1 Å and 3.7 Å, respectively (Supplementary Fig. 10).

For the sample of NBD^WT^-AMPPNP, a total of 720 micrographs were recorded with 200 kV FEI Titan Krios (Gatan K2 camera) at a magnification of 36,000X and at calibrated pixel sizes of 1.157 Å. 32 frames with exposure time of 8 s were collected with defocus ranged from -1.2 to -2.0 μm under low-dose condition. After 2D classification, both hexameric rings and heptameric rings of top views were observed (Supplementary Fig. 9).

Data collection and processing were summarized in S1 Table.

### Crystallization, data collection, and structure determination of the ANK domain

Plate-shaped crystals of CLPB-ANK were obtained within 3 weeks by sitting drop vapor diffusion method at 16°C. Typically, a volume of 1 μL of protein solution at a concentration of 6-12 mg/mL in the GF buffer (5 mM Tris-HCl, pH 7.5, 100 mM NaCl) was mixed with an equal volume of a precipitant well solution of 0.2 M Ammonium citrate tribasic, pH 7.0, 20% (w/v) Polyethylene glycol 3,350. Crystals were directly frozen and stored in liquid nitrogen prior to data collection. The final data set was collected at the Shanghai Synchrotron Radiation Facility (SSRF) beamline BL18U1 (wavelength = 0.97915 Å, temperature = 100K). A 900 diffraction images were collected with oscillation step of 0.2°. The data were merged and scaled using XDS (Kabsch, 2010) and Aimless (Evans, 2011). The initial structure solution was obtained using the molecular replacement program PHASER (McCoy et al., 2007) followed by AutoBuild (Terwilliger et al., 2008) with a prediction structure of CLPB by alpha- fold (Jumper et al., 2021). Further model building was done using Coot and Phenix (Adams et al., 2010; Emsley and Cowtan, 2004), respectively. Data collection and refinement statistics are summarized in (S2 Table) and the atomic coordinates and structure factors have been deposited in the protein data bank (PDB ID: 7XC5).

### Model building and refinement

Model building was based on the prediction structure of CLPB by AlphaFold (Jumper et al., 2021) and the crystal structure of the ANK domain. The models were first docked into the cryo-EM density map using UCSF Chimera (Pettersen et al., 2004), rebuilt manually in Coot and refined (real-space) using Phenix (Adams et al., 2010; Emsley and Cowtan, 2004). Figures preparation and structure analysis were performed with and UCSF ChimeraX (Pettersen et al., 2021) and Chimera (Pettersen et al., 2004).

### Data availability

The cryo-EM density maps have been deposited in the Electron Microscopy Data Bank with the accession codes EMDB-33105, EMDB-33106, EMDB-33109, EMDB-33110 and EMDB-33104 for the CLPB, CLPB^E425Q^, NBD^E425Q^-hexamer, NBD^E425Q^-heptamer and NBD^E425Q^-nanomer, respectively. The atomic model of NBD^E425Q^- nanomer has been deposited in Protein Data Bank with accession code PDB 7XBK.

The coordinate and the structure factor of CLPB-ANK have been deposited in the Protein Data Bank with accession code 7XC5.

## Acknowledgments

We thank the Core Facilities at the School of Life Sciences, Peking University for help with negative staining EM; the Electron Microscopy Laboratory and Cryo-EM Platform for help with data collection; the High-performance Computing Platform for help with computation; the National Centre for Protein Sciences at Peking University for assistance with mass spectrum. The work was supported by the National Science Foundation of China (31725007 to N.G., 31922036 to N.L.), the National Key Research and Development Program of China (2019YFA0508904 to N.G.), and the Qidong-SLS Innovation Fund to N.G.

## Author contributions

D.W. prepared the protein samples (with the help of Y.D.), collected the cryo-EM data, performed EM analysis (with the help of G.W., Y.C., and N.G.), and carried out functional experiments. N.G. and D.W. performed model building and wrote the manuscript. D.W., Y.L., G.L. and J.L. purified the ANK domain and performed the crystallization, data collection, and structure determination.

## Competing interests

Authors declare no competing interests.

Correspondence and requests for materials should be addressed to N.G. and J.L.

---

**S1.**
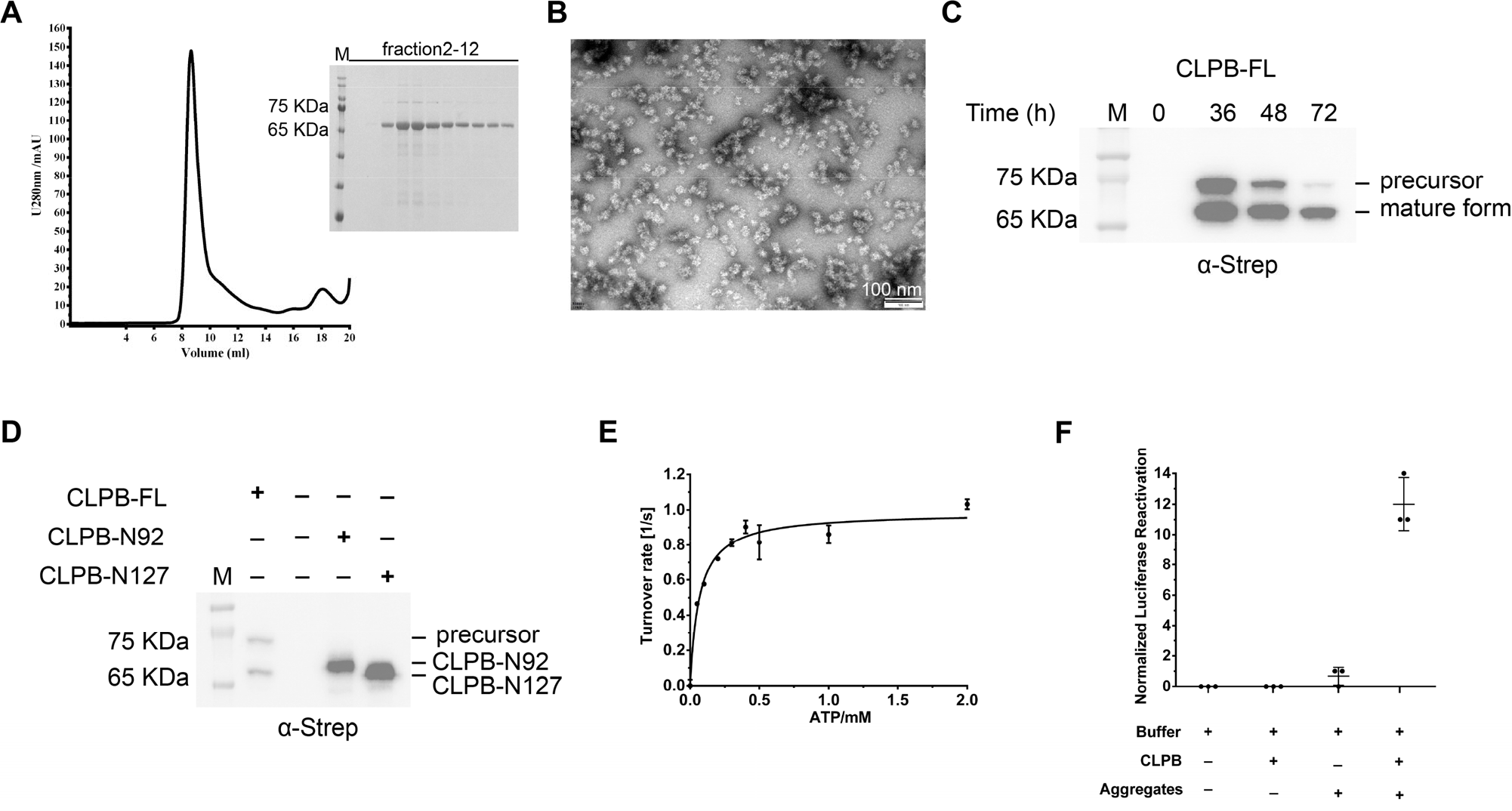
Biochemical and functional characterization of CLPB proteins. **A** The purification of CLPB-N92, analyzing by size-exclusion chromatography (left) and SDS-PAGE (right). **B** Negative staining electron microscopy (nsEM) of the peak fraction in (A). Result shows that CLPB-N92 formed large aggregates and was highly heterogenous in size. **C** Time course of the C-terminal Step-tagged full-length CLPB (CLPB-FL) expression in transiently transfected HEK-293T cells. **D** CLPB-FL, CLPB-N92 and CLPB-N127 expression in transiently transfected HEK-293T cells. Data show that the CLPB-N127 has the same molecular weight as the mature form of CLPB. **E** ATPase activity of CLPB-N127. **F** Disaggregase activity of CLPB-N127.

**S2.**
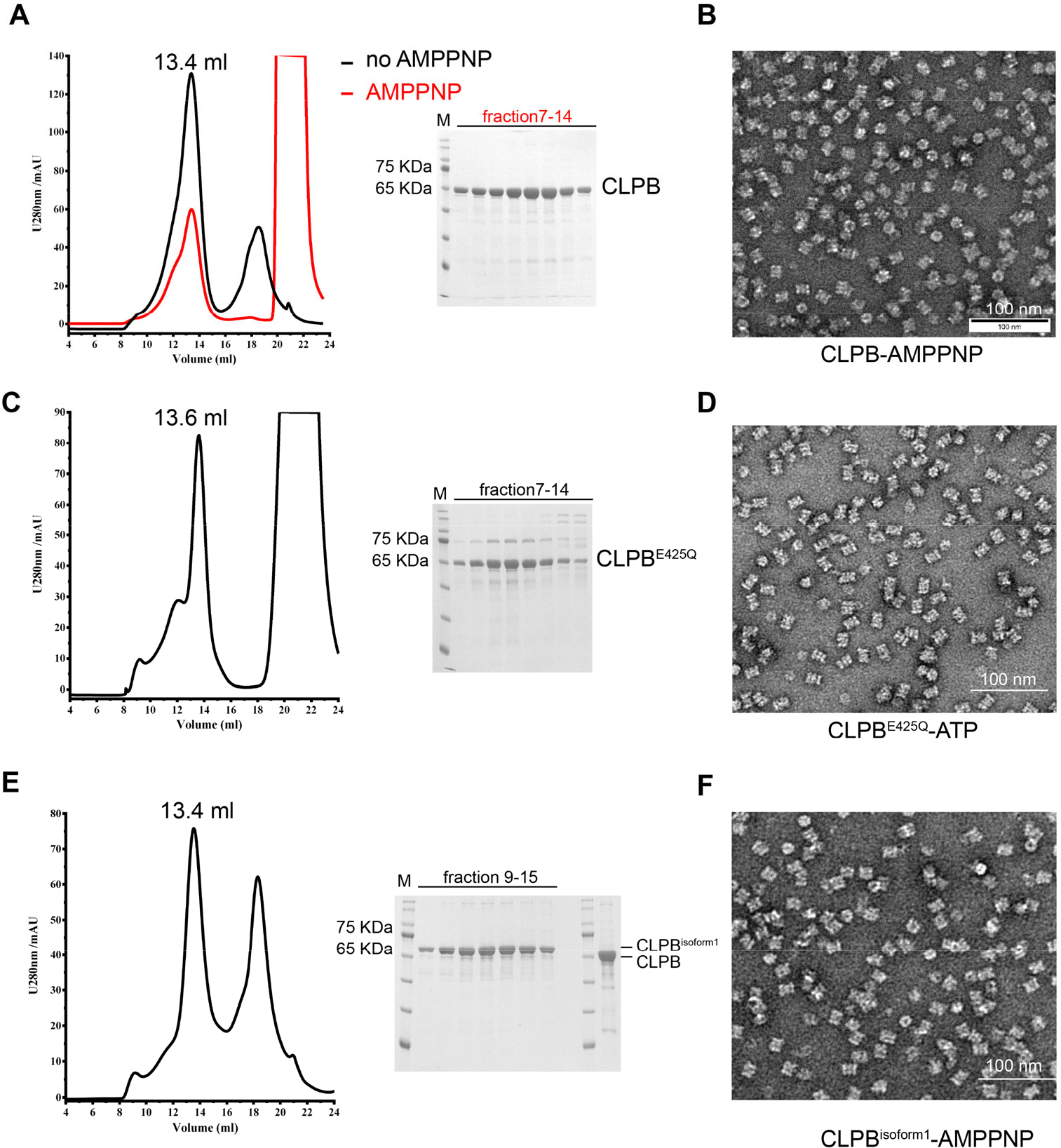
Sample preparation of CLPB-AMPPNP, CLPB^E425Q^-ATP and CLPB^isoform1^-AMPPNP. **A** Purification of CLPB using size-exclusion chromatography with AMPPNP (red line) or without AMPPNP (black line). Corresponding fractions were analyzed by SDS-PAGE (right panel). **B** Representative nsEM image of the peak fraction in (A). **C** Purification of CLPB^E425Q^ using size-exclusion chromatography in the presence of ATP. Corresponding fractions were analyzed by SDS-PAGE (right panel). **D** Representative nsEM image of the peak fraction in (C). **E** Purification of CLPB^isodorm1^ using size-exclusion chromatography. Corresponding fractions were analyzed by SDS-PAGE (right panel). **F** Representative nsEM image of the peak fraction in (E)

**S3.**
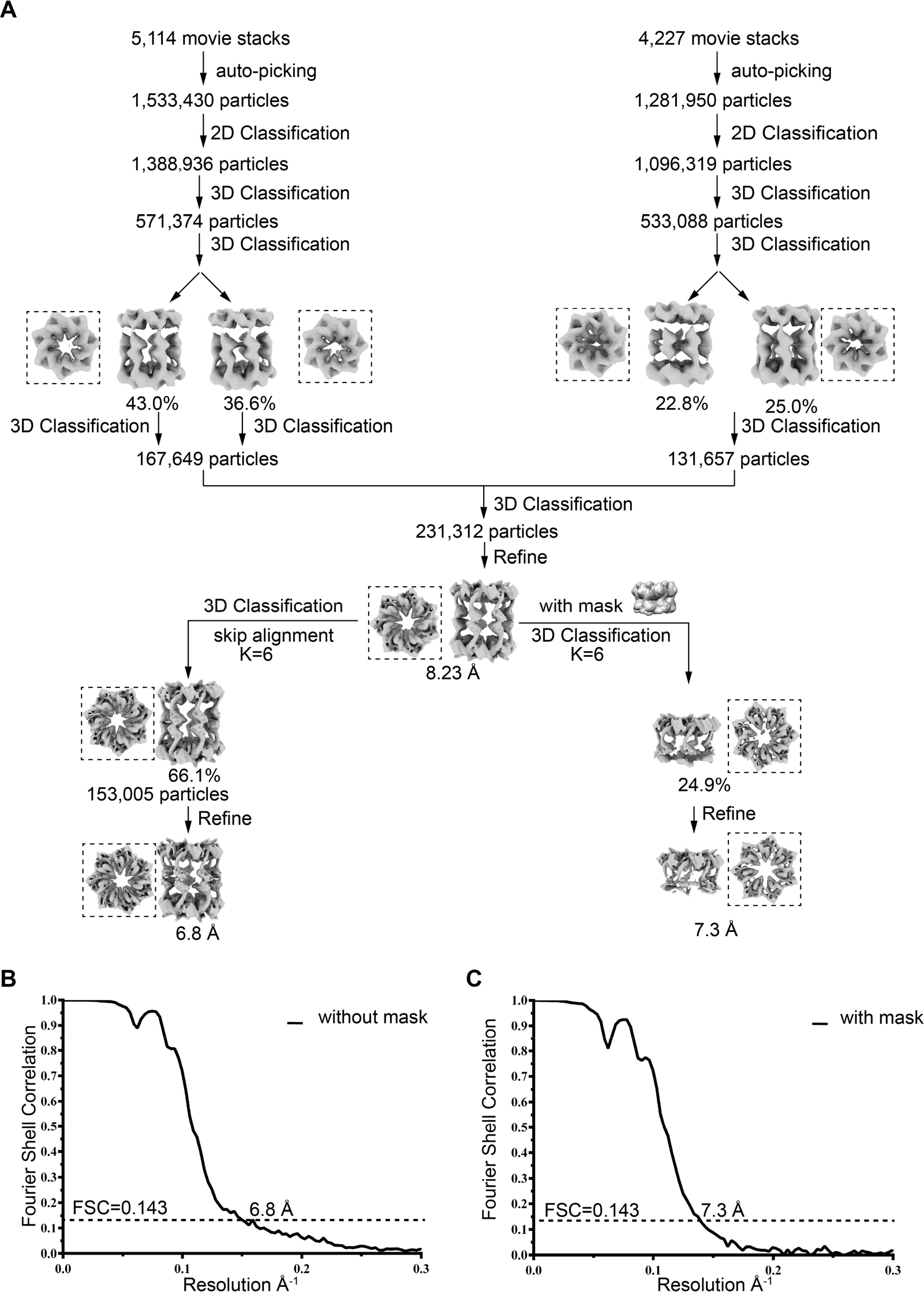
Image processing workflow of the CLPB-AMPPNP dataset. **A** Image processing workflow of the CLPB dataset (see Methods for details). **B, C** Fourier Shell Correlation (FSC) curves for the final cryo-EM map of the double-heptameric complex (B) or heptameric complex (C) using the gold standard FSC 0.143 criteria.

**S4.**
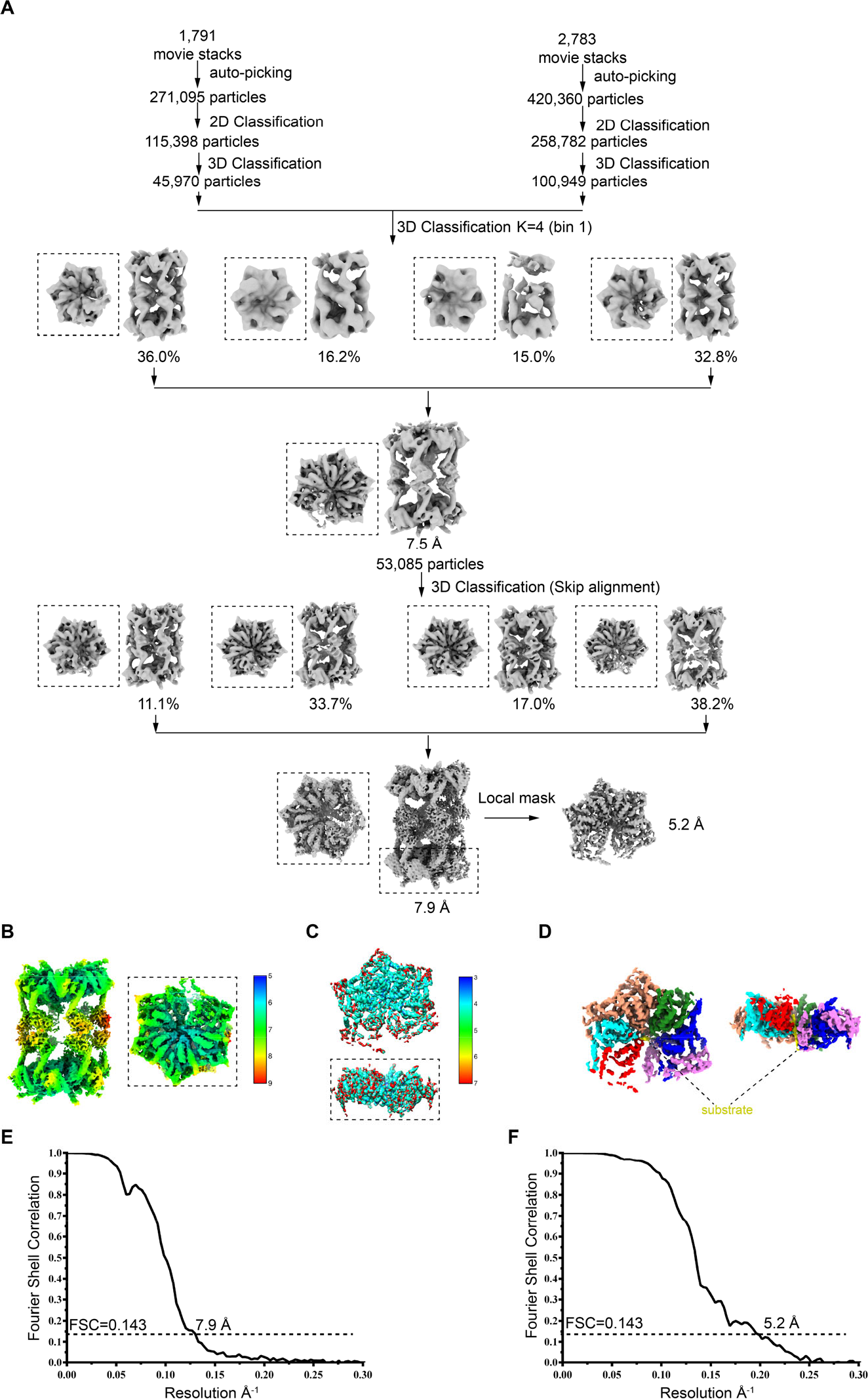
Image processing workflow of the CLPB^E425Q^-ATP dataset. **A.** Image processing workflow of the CLPB^E425Q^ dataset. **B, C** Local resolution estimation of CLPB^E425Q^ double-hexamer (B) or NBD alone (C). **D** Density map of the NBD alone. The central substrate is colored yellow. **E, F** Fourier shell correlation (FSC) curve of the final map of the CLPB^E425Q^ double-hexamer (E) and NBD (F).

**S5.**
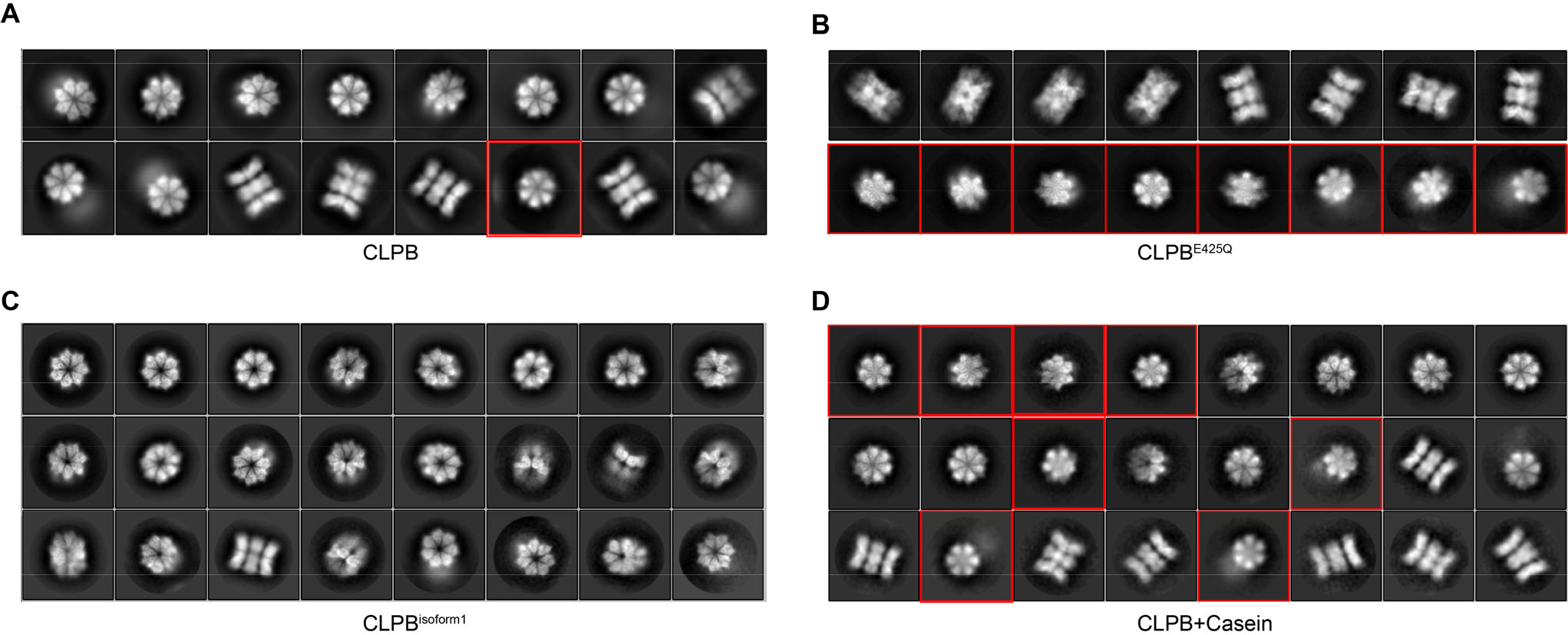
Conversion of the CLPB double-heptamer to double-hexamer upon substrate binding. **A-D** Representative 2D classification averages of CLPB (A), CLPB^E425Q^ (B), CLPB^isoform1^ (C) and CLPB+Casein (D) datasets. The top views with hexameric features are indicated by red boxes.

**S6.**
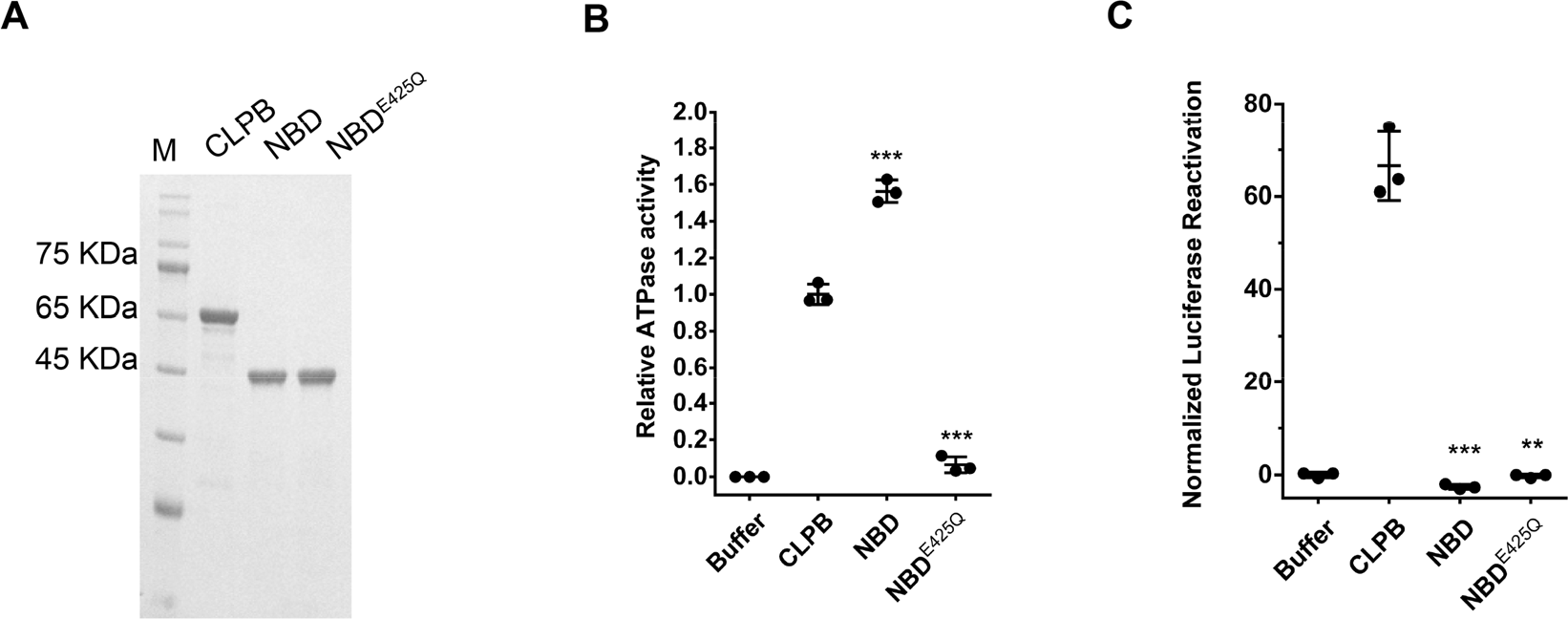
The ANK domain is essential for the disaggregase activity of CLPB. **A** SDS-PAGE analysis of the purified proteins. **B** ATPase assays of CLPB, NBD and NBD^E425Q^. Results show that NBD^E425Q^ has nearly no ATPase activity. NBD has strong ATPase activity. ATPase activity was compared to CLPB (N=3, individual data points shown as dots, bars show mean ± SD, *p<0.05, **p<0.01, ***p<0.0001). **C** Disaggregase activity assay of CLPB, NBD and NBD^E425Q^. The results show that NBD and NBD^E425Q^ abolish the disaggregase activity of CLPB. Disaggregase activity was compared to CLPB (N=3, individual data points shown as dots, bars show mean ± SD, **p<0.01, ***p<0.0001).

**S7.**
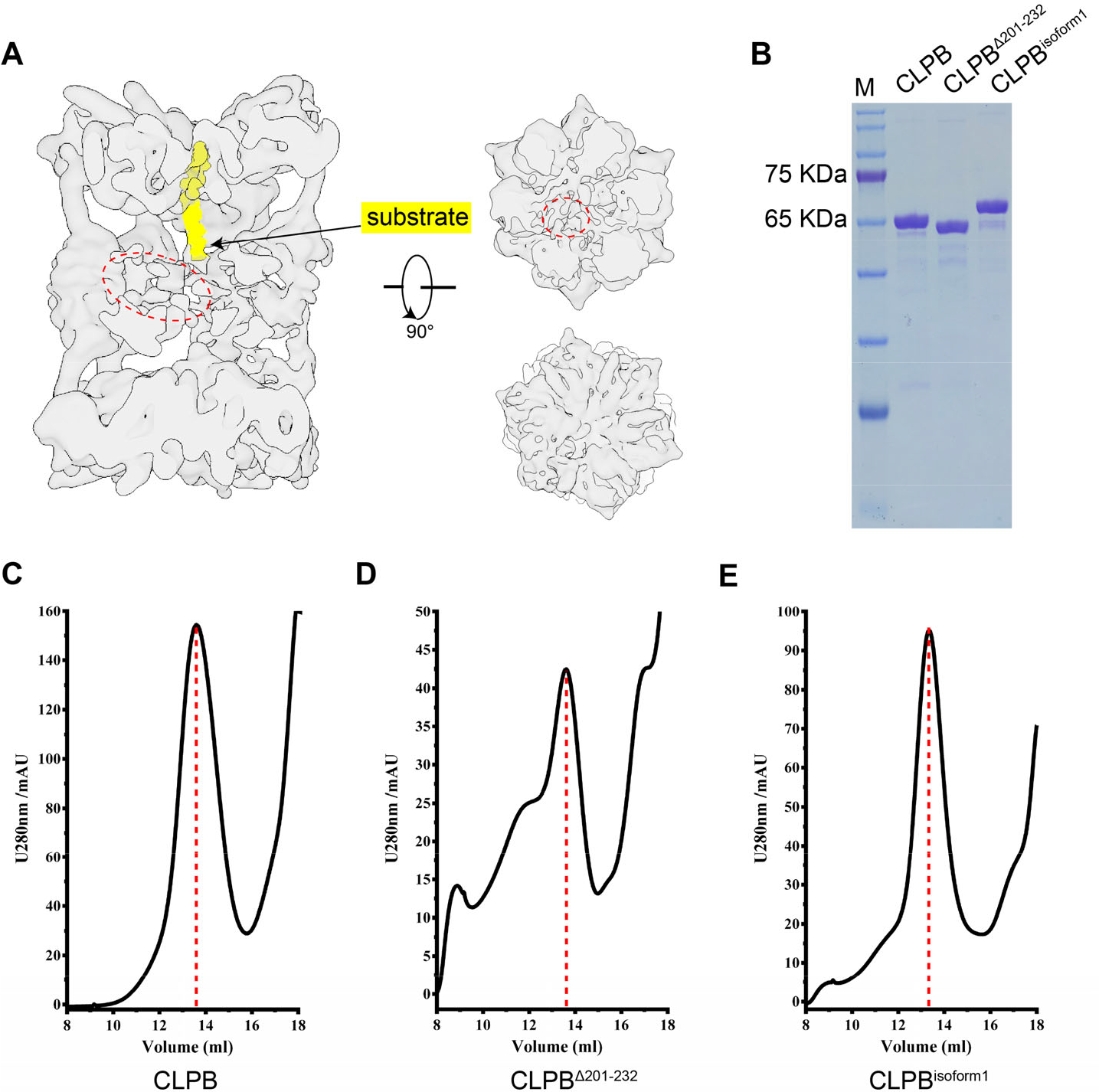
The unique insertion in ANK domain mediates higher-order structure formation and directly contact the substrate. **A** Residual densities around the ANK domains, extending towards the central channel of the CLPB^E425Q^ complex in the substrate-bound state. The extra densities are highlighted by a dotted ellipse. The density of the substrate is shown in yellow. **B** SDS-PAGE analysis of the purified proteins. **C-E** Purification of CLPB, CLPB^Δ201-232^ and CLPB^isoform1^ complexes using size-exclusion chromatography.

**S8.**
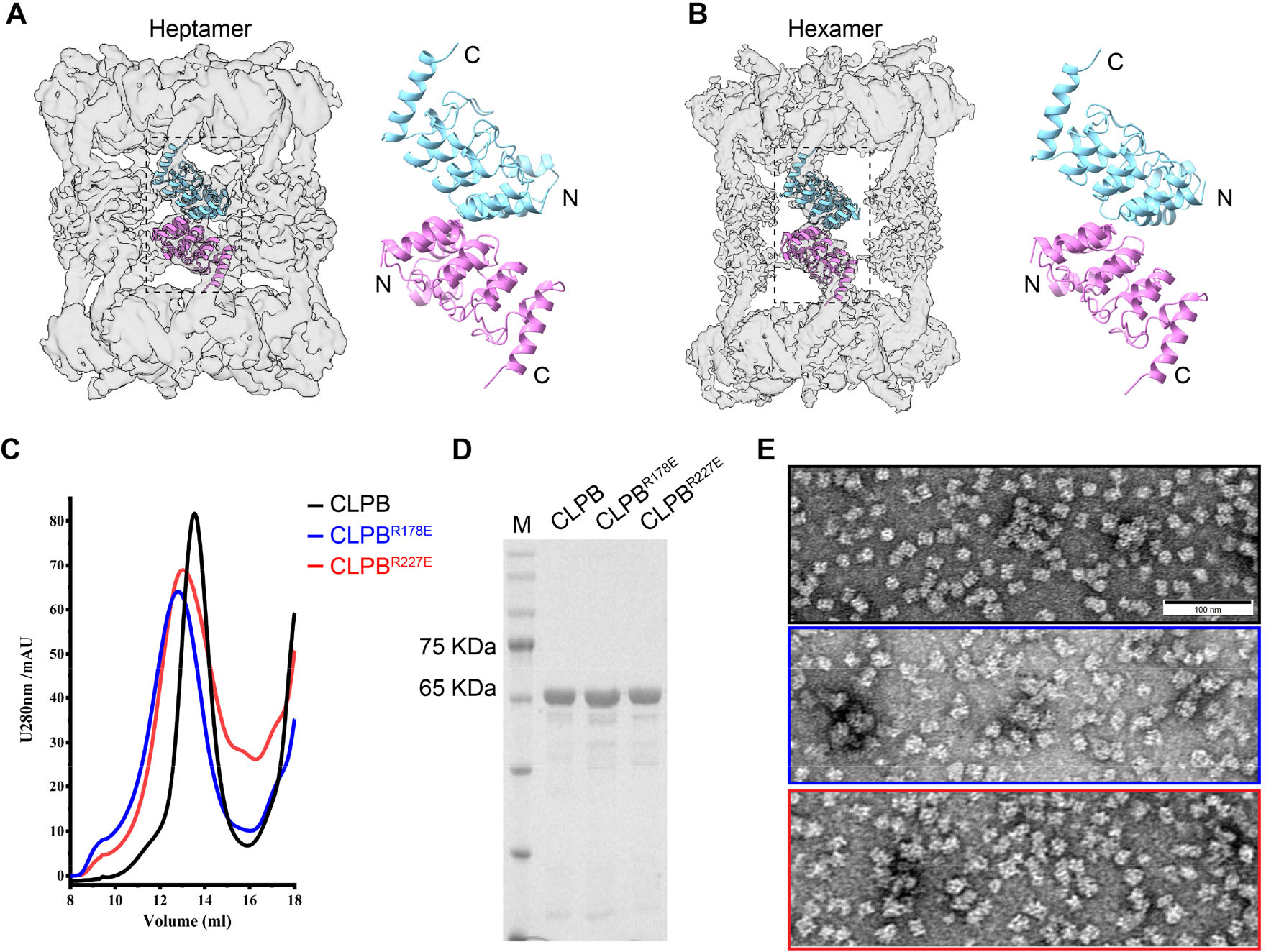
The ANK domain dimer interface in the double-heptamer and double-hexamer. **A, B** Rigid-body fitting of the crystal structure of the ANK domain into the density maps of the double-heptamer (A) and double-hexamer (B). **C** Purification of the CLPB, CLPB^R178E^ and CLPB^R227E^ complexes using size-exclusion chromatography. **D** SDS-PAGE analysis of the purified proteins. **E** Representative nsEM images of the CLPB (black line rectangle), CLPB^R178E^ (blue line rectangle) and CLPB^R227E^ complexes (red line rectangle).

**S9.**
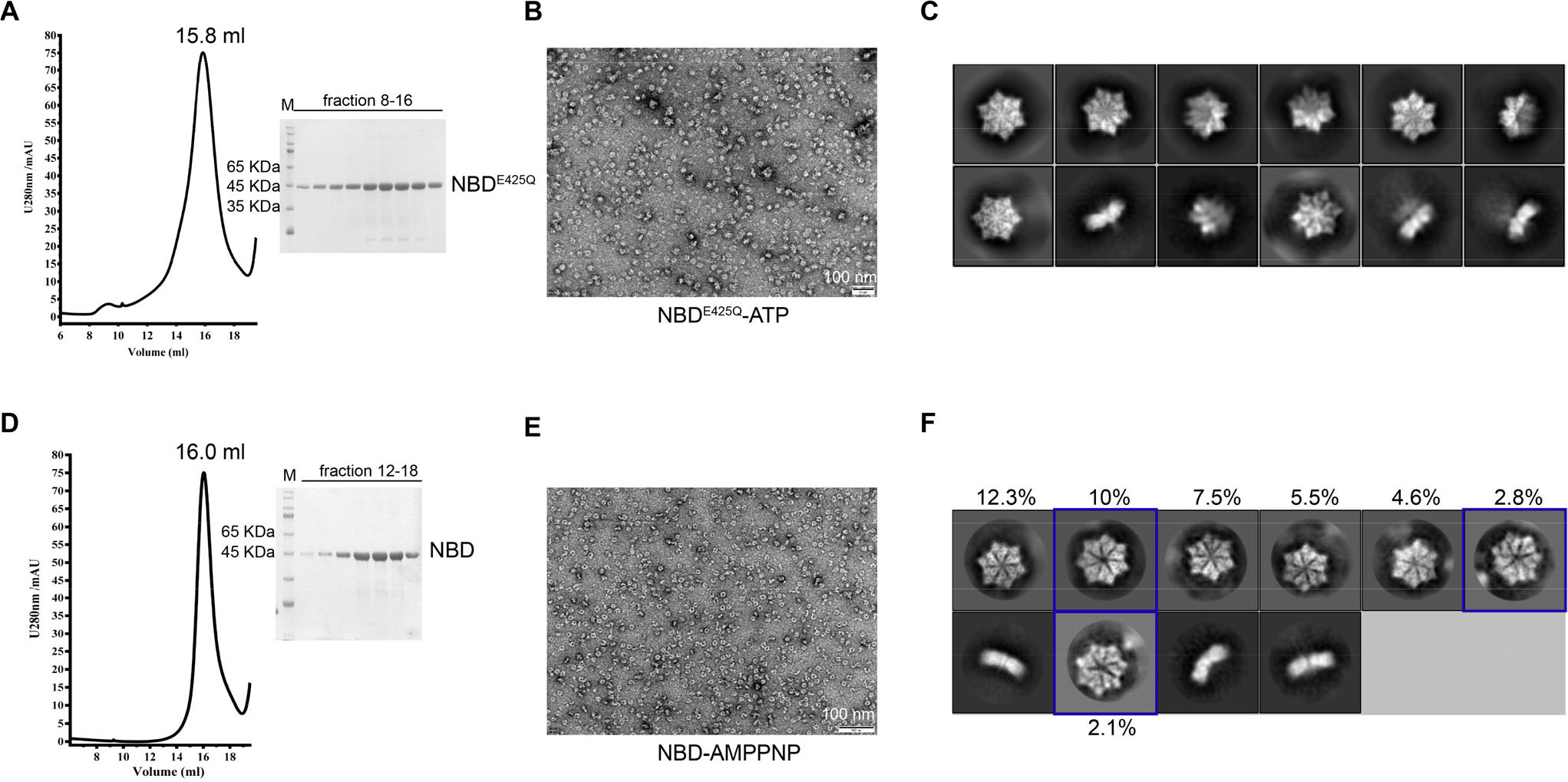
Sample preparation of the NBD^E425Q^-ATPand NBD-AMPPNP complex. **A** Purification of NBD^E425Q^ using size-exclusion chromatography in the presence of ATP. Corresponding fractions were analyzed by SDS-PAGE **B, C** Representative nsEM image (B) and cryo-EM 2D classification averages (C) of the peak fraction in (A). **D** Purification of NBD using size-exclusion chromatography in the presence of AMPPNP. Corresponding fractions were analyzed by SDS-PAGE (right panel). **E, F** Representative nsEM image (E) and cryo-EM 2D classification averages (F) of the peak fraction in (D). The hexameric ring is more compact than hepatameric ring, and the diameter of central pore of hexamer is much smaller than that of heptamer.

**S10.**
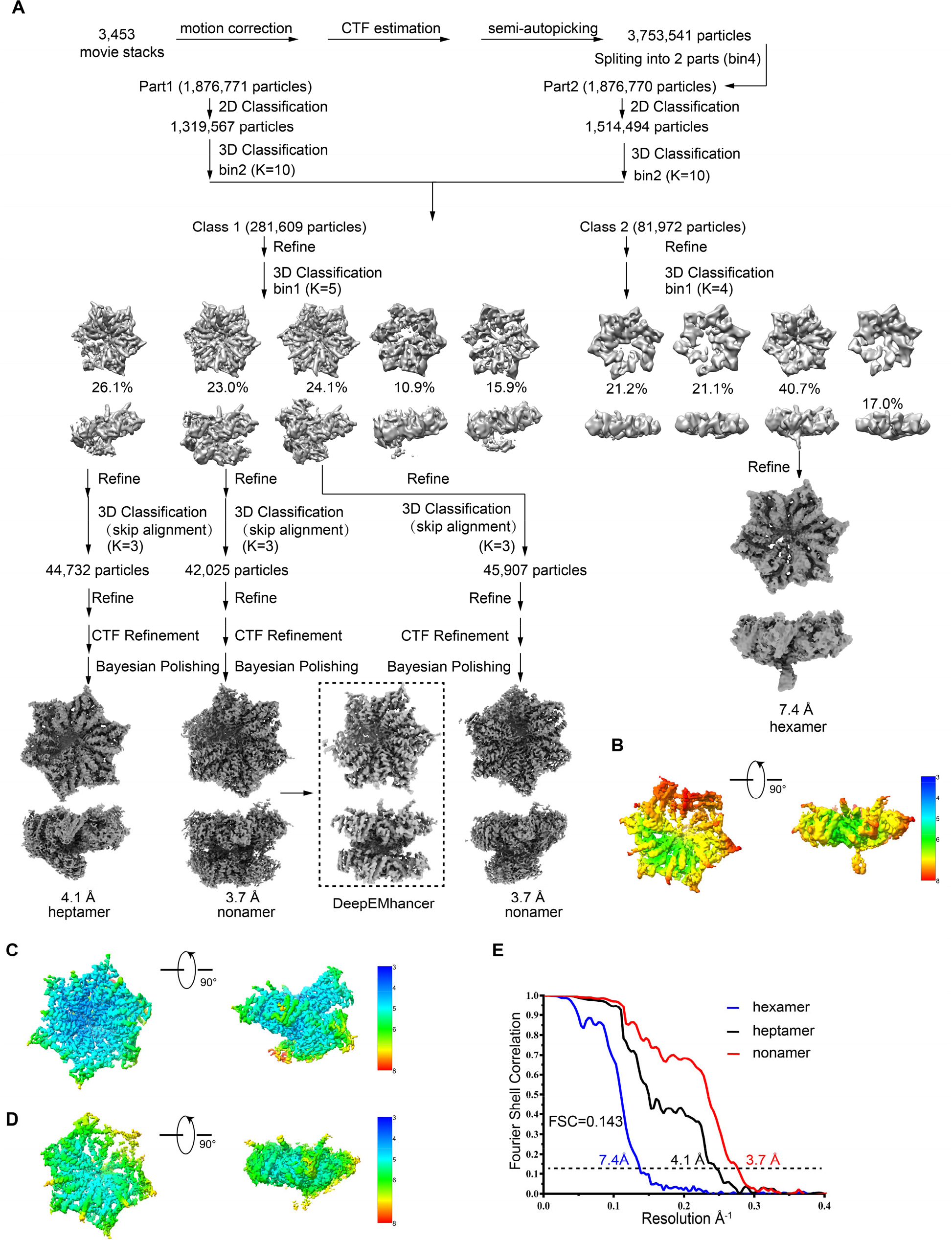
Image processing workflow of the NBD^E425Q^ dataset. **A** Image processing workflow of the NBD^E425Q^ dataset. The micrographs were subjected to motion correction and CTF estimation. The auto-picked particles were subjected to multiple rounds of 2D and 3D classifications. Three different oligomeric arrangements, hexameric, heptameric and nonameric were identified. **B-D** Local resolution estimation of the density maps of the hexamer (B), heptamer (C) and nonamer (D) in (A). **E** Fourier shell correlation (FSC) curves of the final cryo-EM maps of hexamer (blue line), heptamer (black line) and nonamer (red line), using the gold standard FSC 0.143 criteria.

**S11.**
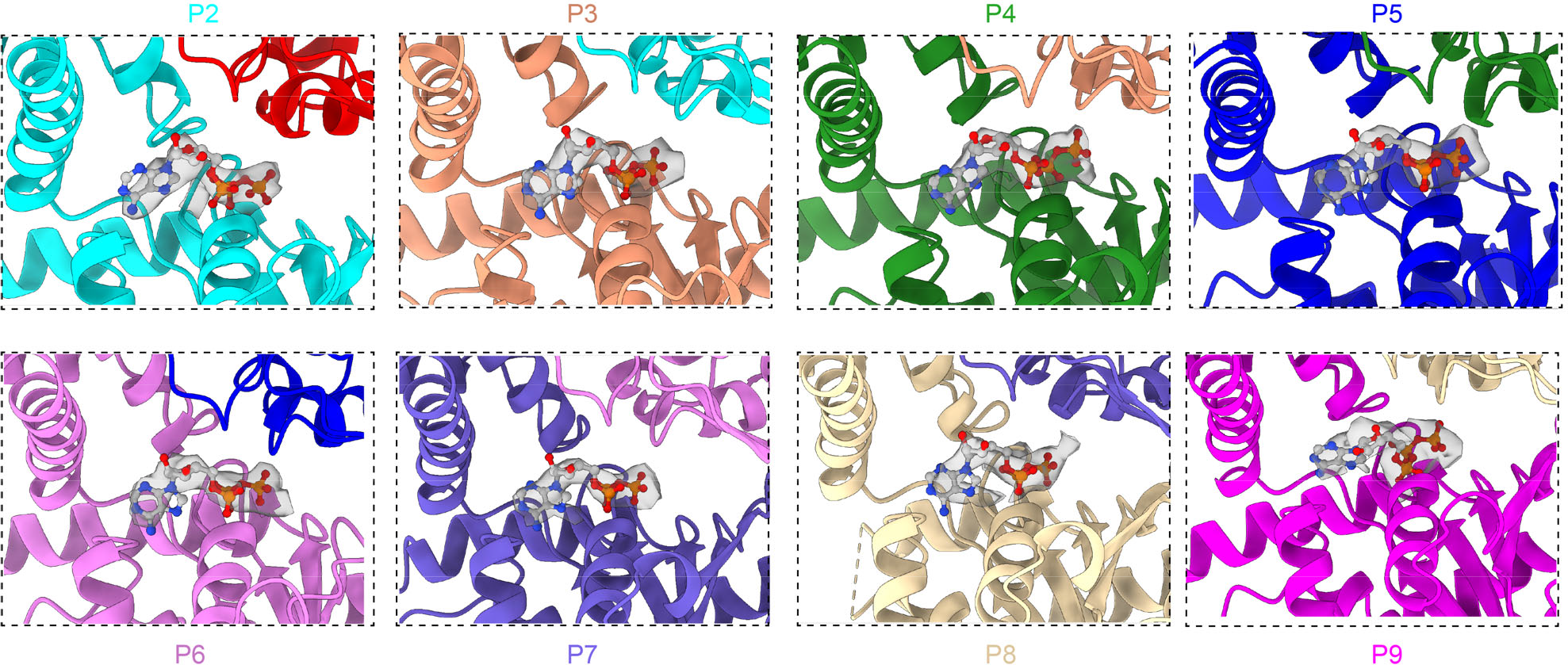
Nucleotide binding states of the nine ATPase sites in the nonamer. ATP occupation of all the ATPase sites in the nonamer. The atomic models are color-coded for different protomers. The segmented density maps of ATP were superimposed with the atomic model.

**S1 Table.** Cryo-EM data collection, refinement and validation statistics.

|  | CLPB | CLPB<sup>E425Q</sup> | NBD<sup>E425Q</sup> |  |  |
| --- | --- | --- | --- | --- | --- |
|  |  |  | Hexamer | Hetamer | Nonamer |
| EMDB/PDB | 33105 | 33106 | 33109 | 33110 | 33104/7XBK |
| **Data collection and processing** |  |  |  |  |  |
| Microscope and camera | Titan Krios K3 | Titan Krios K2 | Titan Krios G3i, K3 |  |  |
| Magnification | 81,000 | 36,000 | 64,000 |  |  |
| Voltage (kV) | 300 | 300 | 300 |  |  |
| Electron exposure (e<sup>-</sup>/Å<sup>2</sup>) | 60 | 58 | 50 |  |  |
| Pixel size (Å) | 1.07 | 1.052 | 1.08 |  |  |
| Map resolution (Å) | 6.8 | 7.9 | 7.4 | 4.1 | 3.7 |
| FSC threshold | 0.143 |  |  |  |  |
| Map resolution range (Å) | 4-7 | 5-9 | 4-8 | 4-8 | 3.5-7 |
| **Refinement** |  |  |  |  |  |
| Map sharpening B factor (Å<sup>2</sup>) | -89.9 | -155.0 | -237.2 | -83.4 | -97.3 |
| **Model composition** |  |  |  |  |  |
| Non-hydrogen atoms | - | - | - | - | 25745 |
| Protein residues | - | - | - | - | 3156 |
| Ligands | - | - | - | - | 8 Mg+8ATP |
| **B factors (Å<sup>2</sup>)** |  |  |  |  |  |
| Protein | - | - | - | - | 88.42 |
| Ligand | - | - | - | - | 66.90 |
| **R.m.s. deviations** |  |  |  |  |  |
| Bond lengths (Å) | - | - | - | - | 0.005 |
| Bond angles (°) | - | - | - | - | 0.863 |
| **Validation** |  |  |  |  |  |
| MolProbity score | - | - | - | - | 1.48 |
| Clash score | - | - | - | - | 5.67 |
| **Ramachandran plot** |  |  |  |  |  |
| Favored (%) | - | - | - | - | 97.04 |
| Allowed (%) | - | - | - | - | 2.93 |

**S2 Table.** Data collection and refinement statistics of the CLPB_ANK.

| CLPB_ANK |  |
| --- | --- |
| **Data collection<sup>a</sup>** |  |
| Space group | C121 |
| Cell dimensions |  |
| *a*, *b*, *c* (Å) | 30.31 67.68 84.98 |
| $\alpha$, $\beta$, $\gamma$ | 90.0 96.05 90.00 |
| Resolution (Å) | 33.84-1.99 (2.04-1.99)<sup>b</sup> |
| *R*<sub>merge</sub> | 0.058 (0.623) |
| *R*<sub>meas</sub> | 0.070 (0.739) |
| *I* / $\sigma I$ | 11.2 (1.9) |
| *CC*(1/2) | 0.998 (0.865) |
| Completeness (%) | 99.4 (99.5) |
| Redundancy | 1.6 (1.7) |
| **Refinement** |  |
| Resolution (Å) | 2.1 |
| No. reflections | 9,966 |
| *R*<sub>work</sub>/*R*<sub>free</sub> (%) | 20.6/23.6 |
| No. atoms |  |
| Protein | 1,287 |
| Water | 29 |
| *B*-factors (Å<sup>2</sup>) |  |
| Protein | 49.0 |
| Water | 43.9 |
| R.m.s. deviations |  |
| Bond lengths (Å) | 0.003 |
| Bond angles (°) | 0.629 |
| Ramachandran stat.(%) | 98.78 / 1.22 / 0.0 / 0.0<sup>c</sup> |

<sup>a</sup> One crystal was used for data collection.
<sup>b</sup> Values in parentheses are for the highest-resolution shell.
<sup>c</sup> Values are in percentage and are for most favoured, additionally allowed, generously allowed, and disallowed regions in Ramachandran plots, respectively.

